# Cartography of opportunistic pathogens and antibiotic resistance genes in a tertiary hospital environment

**DOI:** 10.1101/644740

**Authors:** Kern Rei Chng, Chenhao Li, Denis Bertrand, Amanda Hui Qi Ng, Junmei Samantha Kwah, Hwee Meng Low, Chengxuan Tong, Maanasa Natrajan, Michael Hongjie Zhang, Licheng Xu, Karrie Kwan Ki Ko, Eliza Xin Pei Ho, Tamar V. Av-Shalom, Jeanette Woon Pei Teo, Chiea Chuen Khor, MetaSUB Consortium, Swaine L. Chen, Christopher E. Mason, Tek N Oon, Kalisvar Marimuthu, Brenda Ang, Niranjan Nagarajan

**Affiliations:** Genome Institute of Singapore, 60 Biopolis Street, #02-01 Genome, Singapore 138672, Singapore; Singapore University of Technology and Design, 8 Somapah Rd, Singapore 487372, Singapore; Department of Microbiology, Singapore General Hospital, Singapore 169856, Singapore; Department of Molecular Pathology, Singapore General Hospital, Singapore 169856, Singapore; Duke-NUS Graduate Medical School, Singapore 169857, Singapore; Department of Laboratory Medicine, National University Hospital, 5 Lower Kent Ridge Road, Main Building I, Singapore 119074, Singapore; Department of Physiology and Biophysics, Weill Cornell Medicine, New York 10065, USA; Institute of Infectious Diseases and Epidemiology, Tan Tock Seng Hospital, 11 Jalan Tan Tock Seng, Singapore 304833, Singapore; National University of Singapore, 21 Lower Kent Ridge Road, Singapore 119077, Singapore

## Abstract

There is growing attention surrounding hospital acquired infections (HAIs) due to high associated healthcare costs, compounded by the scourge of widespread multi-antibiotic resistance. Although hospital environment disinfection is well acknowledged to be key for infection control, an understanding of colonization patterns and resistome profiles of environment-dwelling microbes is currently lacking. We report the first extensive genomic characterization of microbiomes (428), common HAI-associated microbes (891) and transmissible drug resistance cassettes (1435) in a tertiary hospital environment based on a 3-timepoint sampling (1 week and >1 year apart) of 179 sites from 45 beds. Deep shotgun metagenomic sequencing unveiled two distinct ecological niches of microbes and antibiotic resistance genes characterized by biofilm-forming and human microbiome influenced environments that display corresponding patterns of divergence over space and time. To study common nosocomial pathogens that were typically present at low abundances, a combination of culture enrichment and long-read nanopore sequencing was used to obtain thousands of high contiguity genomes (2347), phage sequences (1693) and closed plasmids (5910), a significant fraction of which (>60%) are not represented in current sequence databases. These high-quality assemblies and metadata enabled a rich characterization of resistance gene combinations, phage diversity, plasmid architectures, and the dynamic nature of hospital environment resistomes and their reservoirs. Phylogenetic analysis identified multidrug resistant strains as being more widely distributed and stably colonizing across hospital sites. Further genomic comparisons with clinical isolates across multiple species supports the hypothesis that multidrug resistant strains can persist in the hospital environment for extended periods (>8 years) to opportunistically infect patients. These findings highlight the importance of characterizing antibiotic resistance reservoirs in the hospital environment and establishes the feasibility of systematic genomic surveys to help target resources more efficiently for preventing HAIs.

## Introduction

The global epidemic of antibiotic resistance has refocused attention on infection prevention and control measures in hospitals^1^. It is estimated that if the spread of antibiotic resistance grows unchecked, it will cause millions of deaths worldwide, with economic impact of >100 trillion dollars by 2050^2^. In general, hospital acquired infections (HAIs) pose a high healthcare burden in both developed and developing countries^3^. In the U.S., approximately one in 25 acute-care hospital patients have active HAIs each day, translating to 721,800 HAIs each year^4^. It is estimated that 11.5% of such patients will die during hospitalisation^4^. The problem of HAIs is further compounded by the global spread of multidrug resistant organisms (MDROs) that complicate infection management, limit therapy options and result in poorer outcomes^5^. The risk of HAIs can be mitigated through good infection prevention practice, with hand hygiene advocated as one of the key strategies to limit spread of micro-organisms between patients and medical staff^6^.

In addition to human-to-human transfer, the inanimate hospital environment is another key node in the transmission network, with mounting evidence that it harbors opportunistic antibiotic resistant pathogens that contribute to HAIs^7,8^. Reinforced environmental cleaning measures have also been shown to be effective in reducing HAIs^9-11^. As a built-environment with unique clinical significance, the microbial ecology and uncharacterized genetic reservoirs of hospital environments are of special interest for both infection epidemiology and microbiology. For example, the transmission and recombination profiles of antibiotic resistance genes (ARGs) in hospital environments remains largely unknown, and could be valuable in gauging the potential risk for emergence of novel resistance combinations in specific species. Similarly, comparative genomics of hospital adapted strains and epidemic strains can help in identifying the source of HAI outbreaks and inform infection control. While large-scale surveillance holds the promise to reveal clinical and biological insights pertaining to the hospital environment microbiome as a functional reservoir of pathogens and ARGs, significant technological challenges still remain. Traditionally, efforts to survey the hospital environment have been focused on culture-based isolation of specific pathogens. Each isolate is then individually characterized via functional profiling, genotyping and/or whole genome sequencing^12-17^. This is a laborious process, prone to isolation bias and precludes insights into overall microbial community structure and interactions with the hospital built-environment to impact the transmission of HAIs^18^.

The development of metagenomics has made it possible to profile the entire community structure, characterizing individual microbes without the need for isolation, and represents a high-throughput and scalable method for surveying the hospital environment microbiome^19^. This capability has been leveraged in the form of 16S rRNA sequencing to study bacterial diversity, particularly in intensive care units (ICU), in several early studies^20-23^. In a landmark study in 2017, Lax et al used this approach to extensively characterize microbial ecology, colonization and succession patterns in a newly built hospital environment (a subset with shotgun metagenomics)^24^. Using bioinformatics approaches, they were also able to identify ecological signatures of exchange of bacteria between the hospital environment, patients and healthcare workers. In general, however, 16S rRNA sequencing precludes the detailed analysis of nosocomial strains, resistomes, metabolic pathways, and transmission of pathogenomes^25^. The use of Illumina shotgun metagenomics by Brooks et al was key to characterizing strain polymorphisms and relatedness of pathogens in low diversity neonatal ICU environments^26^. Several limitations remain for the use of shotgun metagenomics in general, including low microbial biomass, the presence of nosocomial strains at low relative abundances, the presence of multiple strains, computational constraints in strain-level analysis^27^, and the shortcomings of short-read metagenomics for assembling high-contiguity, strain-resolved genomes for detailed genetic analysis^28^.

The availability of portable, real-time, long-read sequencing platforms presents new opportunities and challenges for pathogenome and resistome monitoring^29,30^. In this study, we combined extensive short-read shotgun metagenomic sequencing of multiple sites, wards and timepoints (428 samples), with enrichment and nanopore sequencing of antibiotic-resistant mixed cultures (1661 samples), to provide the most extensive genetic characterization of hospital environments to date. The combination of metagenomic surveys (short-read based) with detailed genomic analysis of nosocomial strains (long-read based) is ideal for studying the interplay between dissemination, abundance and turnover patterns of pathogens and ARGs with the abiotic components of the hospital environment, informing the development of targeted cleaning protocols and prioritization of high-risk areas for cleaning. Nanopore metagenomics enabled us to generate thousands of high-contiguity genomes (2347), phage sequences (1693) and closed plasmids (5910), unveiling the significant under-characterized genetic diversity (>60% novel) for potentially pathogenic microbes. These were harbored in two distinct ecological niches characterized by biofilm-forming and human microbiome associated bacteria, with divergent patterns of temporal and spatial variation. The availability of a large collection of high-quality genomes helped establish through phylogenetic analysis the notable observation that multi-drug resistant strains are more likely to be widely distributed and stably colonizing across hospital sites. In addition, the genomes delineated common resistance gene combinations, phage diversity and plasmid architectures, revealing the dynamic nature of hospital environment resistomes and their understudied reservoirs. Further genomic comparisons with clinical isolates across multiple species supports the hypothesis that multidrug resistant strains can persist in the hospital environment for extended periods of time (>8 years) to opportunistically infect patients. Together, these findings highlight the importance of characterizing hospital environment microbiomes for understanding its niches and genetic reservoirs, the need for improved methods of disinfection, as well as the feasibility and cost-effectiveness of large-scale genomic surveys for informing infection control measures.

## Results

### Hospital environment microbiomes offer distinct ecological niches for opportunistic pathogens and ARGs

A diverse set of sites (n=7) of concern for infection control^31-33^, and different room types distributed around the building (5 single-bed isolation rooms, 4 MDRO and 4 standard 5-bedded wards) were selected for initial 2-timepoint sampling (1 week apart) of a tertiary care hospital in Singapore (45 beds, 4% of total; 179 sites, 358 samples; **Fig. 1a** and **Suppl. File 1**). Illumina shotgun metagenomic sequencing (2×101bp) was used to deeply characterize each sample (30 million reads on average; 3/358 libraries excluded due to low biomass) and obtain taxonomic and functional profiles, as well as resistomes (**Fig. 1b, Suppl. File 2**; **Methods**). Controls, spike-ins and validation experiments were used to assess and account for the impact of kit contaminants on low biomass samples^34^, with likely contaminant species being identified using batch and correlation analysis^34^, and filtered from profiles (**Suppl. Note 1, Suppl. File 2**; **Methods**). Taxonomic profiles were visualized using a principal coordinates analysis (PCoA) plot to identify two distinct microbial community configurations in the hospital environment (CTA and CTB; **Fig. 2a**). While community type A (CTA) sites were more taxonomically diverse (Wilcoxon p-value<10^−3^; **Suppl. Fig. S1**), and largely high-touch surfaces with frequent contact from patients and health-care workers^35^, community type B (CTB) represents sites that are increasingly of concern for infection control for their propensity to harbor MDROs^16,32,36^. Joint analysis of these community types helped identify key taxonomic features that differentiate them, including several human microbiome-associated (e.g. *Cutibacterium*) and aquatic/terrestrial environment associated (e.g. *Achromobacter*) genera in community types A and B respectively, though not all genera (e.g. *Pseudomona, Acinetobacte*r and *Ralstonia*) could be defined in these terms (**Fig. 2bi**). At the species level, key differences observed include the enrichment of common skin bacteria (e.g. *Cutibacterium acnes* and *Staphylococcus epidermidis*) and common biofilm-associated organisms in hospitals (e.g. *Elizabethkingia anophelis* and *Serratia marcescens*) in CTA and CTB sites respectively, though their occurrences were not mutually exclusive, indicating shared influences (**Fig. 2bii**). Comparison of hospital microbiome CTA and CTB sites, with similar sites in an indoor office environment (n=30, Office; **Suppl. File 1, Methods**) and other high-touch environmental microbiomes^37^ (n=99, MetaSUB Singapore; **Suppl. File 1**), further highlighted the distinctness of hospital environments and their community types (**Suppl. Fig. S2**), and corresponding utility as an organizing principle for studying clinical impact^22,38^.

**Figure 1:**
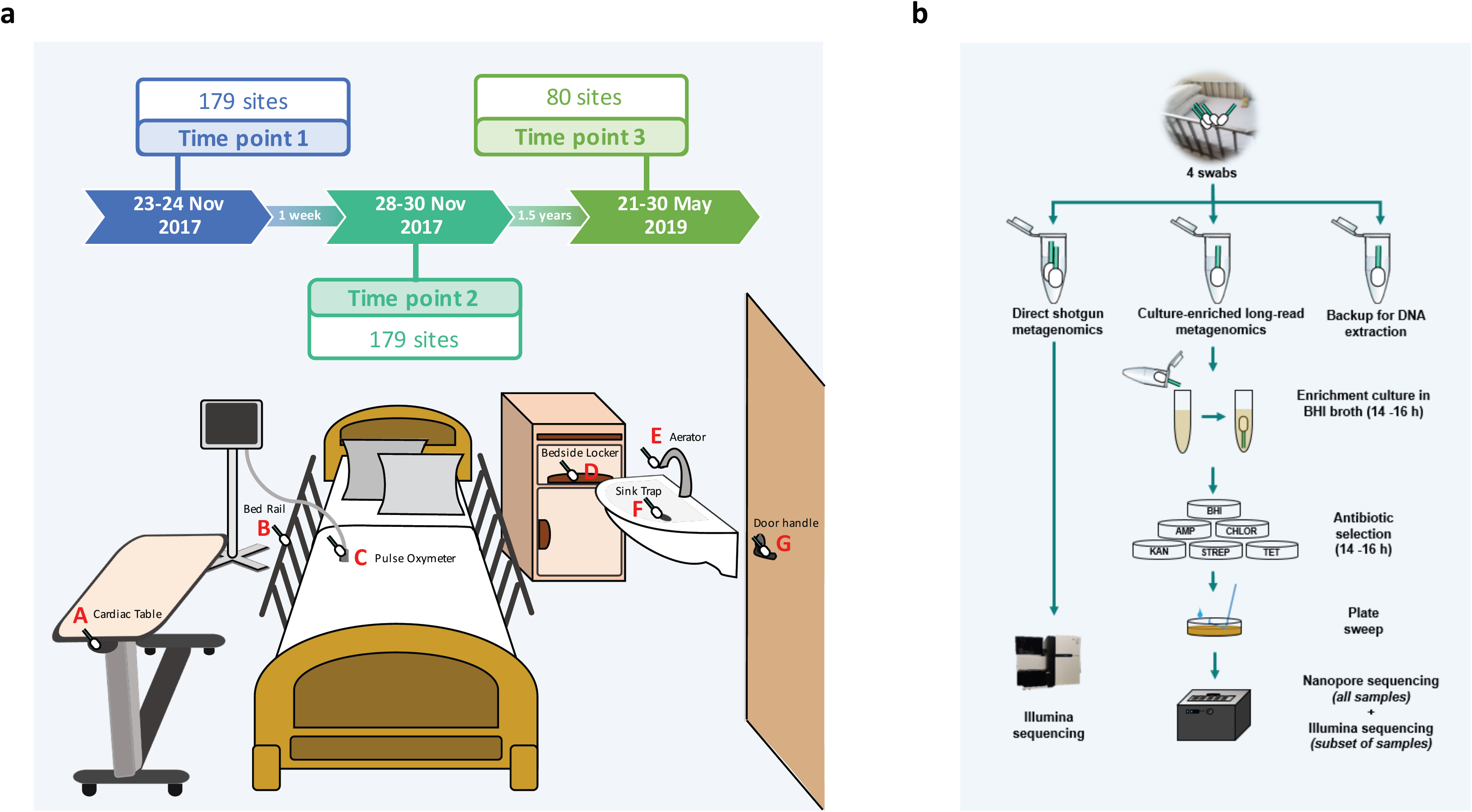
Overview of sampling sites and analysis workflow. (a) Diagram showing the various sites that were sampled, including cardiac tables (A), bedrails (B), pulse oxymeters (C), bedside lockers (D), sink aerators (E), sink traps (F) and door handles (G). Each ward (MDRO and Standard) had five beds (sites A-D individually sampled) and one sink (E, F) and no doors, while isolation rooms had one bed (A-D), one sink (E, F) and a door (G). (b) Diagram showing the analysis workflow for the four swabs that were collected from each site in terms of culturing, DNA extraction and sequencing. Samples from each site were analyzed with shotgun metagenomic sequencing on the Illumina platform and multiple (n=6) culture-enriched quasi-metagenomics on a GridION system.

**Figure 2:**
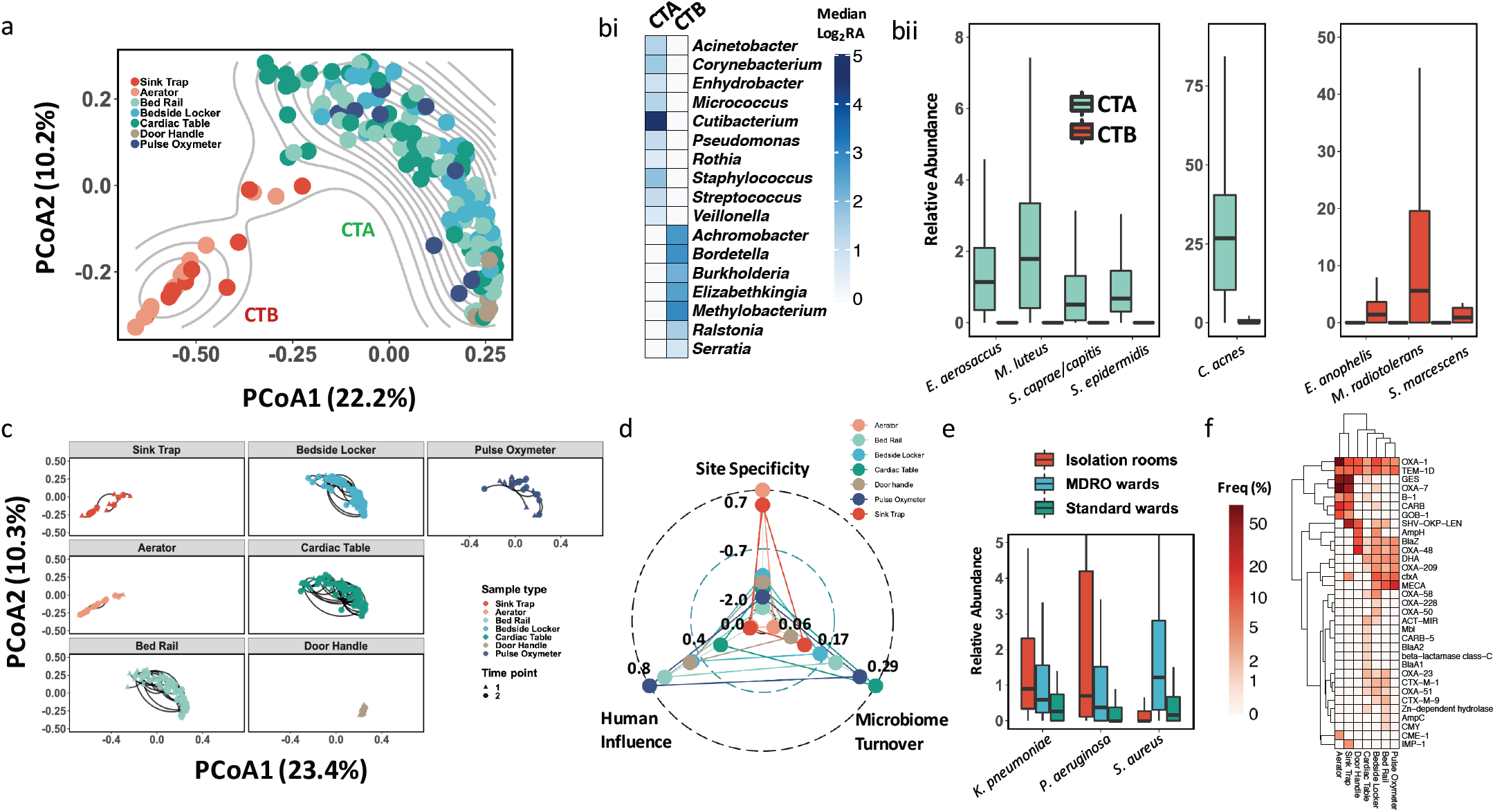
Distinct ecological niches in the hospital environment for microbes and ARGs. (a) Principle coordinates analysis plot based on genus-level Bray-Curtis dissimilarity of taxonomic profiles indicating two distinct community types (denoted as CTA and CTB) for microbiomes from the hospital environment. (bi) Heatmap showing relative abundances (log-scale, Log_2_RA) for differentially abundant genera between community types CTA and CTB (FDR-adjusted Wilcoxon p-value<0.01) (bii) Boxplots showing relative abundances for differentially abundant species between community types CTA and CTB (FDR-adjusted Wilcoxon p-value<0.01). (c) Principle coordinates analysis plot (genus-level Bray-Curtis dissimilarity) showing how much environmental microbiomes vary over time (lines connect two timepoints, 1 week apart) for different sites. (d) Radar plot showing the microbiome turnover index (fraction of taxa that are gained or lost across timepoints), human influence index (fraction of human reads) and site specificity index (uniqueness of site-specific taxonomic composition in relation to physically proximal sites). A positive site specificity index indicates a stronger site-specific microbiome composition signature. (e) Boxplots showing relative abundances of common nosocomial pathogens that were differentially abundant across ward types in high human-contact sites (FDR-adjusted Kruskal-Wallis p-value<0.01). (f) Heatmap depicting the frequency of detection for beta-lactamases at different sites in hospital wards. Multiple carbapenemases and the mecA gene were detected as part of the resistomes that were primarily defined by the community types (CTA and CTB). Boxplots are represented with center line: median; box limits: upper and lower quartiles; whiskers: 1.5× interquartile range; outlier points not included in visualization.

Microbiomes associated with the two community types exhibited varying stability across the two sampled timepoints, with CTA sites demonstrating larger fluctuations (except door handles; Wilcoxon p-value<10^−3^; **Fig. 2c**). In general, microbial profiles diverged with physical distance (within bed, within wards and across wards) and time (1 week apart), with temporal variability within a week being lower than spatial variability within a ward (Wilcoxon p-value<10^−3^; **Suppl. Fig. S3a**). Analysis of a subset of sites (n=80) resampled at a third timepoint >1 year after the first two timepoints, confirmed the long-term stability of community types for various sites (**Suppl. Fig. S3b**). The microbial composition of any particular site is however expected to be influenced by a range of factors including abiotic conditions (humidity, temperature, surface type), seeding from microbial reservoirs (human or environmental) and exchange across sites. Based on the sequencing data, we computed various scores to quantify these factors including a *microbiome turnover* index (fraction of taxa that are gained or lost across timepoints), *human influence* index (fraction of human reads) and *site specificity* index (uniqueness of site-specific taxonomic composition in relation to physically proximal sites), each of which exhibits significantly correlated trends across the sampled timepoints (**Suppl. Fig. S4a**). The computed indices reinforce the notion that CTB sites (primarily sink traps/aerators) tend to have stable compositions (low turnover) based on site-specific biofilm configurations with limited human microbiome seeding (**Fig. 2d**). Sites associated with community type A tend to have higher human influence (Wilcoxon p-value<10^−15^) and microbiome turnover indices (Wilcoxon p-value<10^−4^), though they are not directly correlated, and tend to have weaker site-specificity (Wilcoxon p-value<10^−12^), concordant with a model where human activities (patient discharge/admittance events) have a systemic role in shaping their compositions (**Fig. 2d**). Species that were enriched in CTA sites were also observed in CTB sites (and *vice versa*), but tended to have higher turnover in these cases (**Suppl. Fig. S4b**), with a few exceptions such as *Siphoviridae* having high turnover in both CTA and CTB sites.

We observed that overall patterns of microbiome variability were consistent across ward types, though isolation rooms exhibited lower variability across two timepoints (**Suppl. Fig. S5**). In line with MDRO management guidelines in Singapore^39^, patients colonized with Carbapenem-resistant Enterobacteriaceae (CRE; e.g. *K. pneumoniae*) were typically warded in single-bed isolation rooms, while methicillin resistant *Staphylococcus aureus* (MRSA) carrying patients were largely assigned to MDRO wards. Analysis of differentially abundant common nosocomial pathogens (curated from https://www.cdc.gov/hai/organisms/organisms.html and published literature^4,40^) across ward types identified *K. pneumoniae* and *S. aureus* as being enriched in CTA sites for isolation rooms and MDRO wards respectively, providing further evidence for the influence of patient microbiomes on CTA sites (**Fig. 2e**). Consistent with the observed taxonomic differences, community types A and B also harbored distinct complements of ARGs in their resistomes (**Fig. 2f, Suppl. Fig. S6**). While some resistance genes were frequently detected in CTB sites (e.g. ges, oxa-7 in **Fig. 2f**), CTA sites carried a wider diversity of genes at lower frequencies. Despite the growing recent focus on CTB sites as drug resistance gene reservoirs^16,36^, some clinically important resistance genes such as oxa-23 (carbapenemase) and mecA (methicillin resistance) were more frequently found in CTA sites, while others such as imp-1 (carbapenemase) and cme-1 (extended-spectrum beta-lactamase) were more common in CTB sites. Different sites also exhibited distinct resistome patterns, e.g. specific tetracycline (tetC) and macrolide (mphE) resistance genes were highly enriched only in aerators, while vancomycin resistance genes were only seen in bedside lockers and bedrails, highlighting the importance of considering site and ward-specific patterns when defining infection control and drug resistance mitigation strategies (**Suppl. Fig. S6**). While CTB sites revealed a higher proportion of ARGs as being consistently detected across timepoints compared to CTA sites (**Suppl. Fig. S7a**), overall hospital environment microbiomes exhibit significantly higher abundance (>3×; compared to MetaSUB Singapore and >12× compared to office sites, Wilcoxon *p*-value<10^−15^ for both comparisons, **Suppl. Fig. S7b**) and diversity (Wilcoxon *p*-value<10^−15^; **Suppl. Fig. S7c**) of ARGs compared to other high-touch urban environmental microbiomes. Even though the presence of ARGs does not always translate to resistance phenotypes, these results further underscore the distinctness of hospital environment microbiomes as ARG reservoirs^42^.

### Quasi-metagenomics with nanopore sequencing reveals distribution of multidrug resistant opportunistic pathogens in the hospital environment

Based on Illumina shotgun metagenomic profiles we noted that common nosocomial pathogens were generally present at very low relative abundances (median relative abundance <0.5%; **Fig. 3a**) in hospital environments (even though this was higher than in other urban sites; **Suppl. Fig. S7d**), precluding detailed genomic characterization and analysis of transmission patterns, resistance gene combinations and plasmids. The distribution of common nosocomial pathogens exhibited site-specific patterns (PERMANOVA p-value<0.001; **Fig. 3a**), in agreement with the distinct ecological niches observed in the hospital environment (**Fig. 2a, b**), and indicated that enrichment cultures could capture a diverse set of species. We exploited this observation to use a culturing, antibiotic selection (5 antibiotics) and metagenomic nanopore sequencing approach (**Fig. 1b**) to obtain a large database of high-contiguity genome assemblies (n=2347) from the hospital environment (median N50 >1Mbp; **Fig. 3b, Suppl. File 3**; **Methods**), expanding substantially on genomic resources established by previous hospital environment surveillance studies^16,17^. Overall, a large percentage of sites led to viable initial cultures (>95%), with antibiotic selection still resulting in growth in >80% of plates (1495/1790), and >42% of sites resulting in cultures for all 5 antibiotics. Control swabs led to no cultures (0/10) confirming that cultures were not likely due to contamination (**Methods**), and further testing of isolates for antibiotic resistance confirmed that the vast majority of strains in the cultures were likely antibiotic resistant (99%; **Suppl. Note 2**).

**Figure 3:**
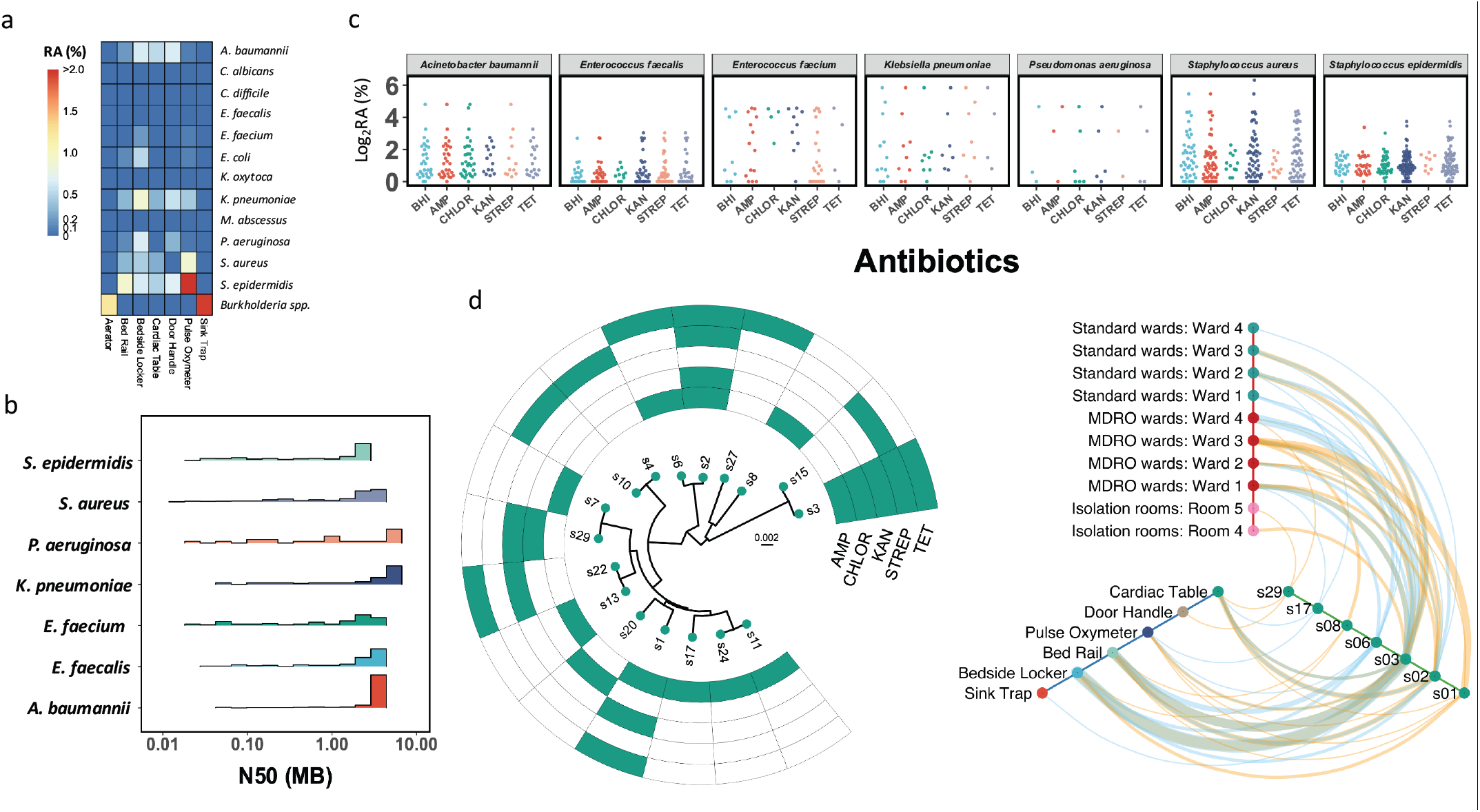
Genome-resolved characterization of nosocomial multidrug resistant strains that spread and persist at low relative abundances in the hospital environment. (a) Heatmap displaying the distinct median relative abundances (RA) of common nosocomial pathogens at different sites in hospital environments (PERMANOVA p-value<0.001). (b) Distribution of assembly contiguity statistics (N50=fragment size s.t. more than 50% of the genome is in longer sequences) for common nosocomial pathogens, highlighting the high genomic contiguity that was obtained (median N50 >1Mbp). (c) Dotplots highlighting that genomes can be rapidly obtained for several nosocomial pathogens despite their low relative abundances in corresponding environmental microbiomes (y-axis), through an enrichment and long-read metagenomic sequencing based protocol. Represented species are associated with more than 20 genome drafts in the overall database of 2347 genomes. (d) (Left panel) Phylogenetic relationships of *S. aureus* derivative clusters (>99.99% ANI; each node represents the consensus genome for the cluster) detected in the hospital environment together with their antibiotic resistance profiles. The scale bar represents the number of substitutions per site in the core alignment. (Right panel) Hive-map representation showing localization of *S. aureus* clusters that spread (detected at 2 or more locations) and/or persist (detected in timepoints 1 and 2) in the hospital environment. Orange lines represent occurrence at time point 1 while blue lines represent occurrence at timepoint 2. Line thickness represents the number of instances of such occurrences. Note that multidrug resistant strains such as s3, s2 and s1 tend to be more widely distributed and persistent in the hospital environment.

DNA was extracted from 1661 plates and sequenced on the GridION system to provide 535Mbp of nanopore sequencing on average per sample (median read length >2.5kbp). Long read metagenomic assembly enabled the frequent reconstruction of megabase-pair sized contigs (versus average N50 <5kbp for Illumina metagenomic assemblies) as the communities were largely simple (**Suppl. Fig. S8, Fig. 3b**; **Methods**). An evaluation of these draft genomes based on evolutionarily conserved single-copy genes confirmed that they were on average of high quality (completeness >99%, contamination <0.5%; **Methods**). In total, we obtained genomes for 69 different known species from the hospital environment, 40% of which belonged to common nosocomial pathogens (**Methods**). Our results confirm the viability of these species in different hospital environments and the ability to substantially enrich them for sequencing and genome reconstruction compared to their acutely low abundances in the hospital environment microbiome (median relative abundance = 0.68%, averaged across species; **Fig. 3c**). Large-scale homology analysis of these genomes with public databases^43-45^ also helped identify 13 (out of >80) species-level clusters (from 11 different genera including *Bacillus, Pseudomonas* and *Staphylococcus*) which do not have representatives from known species, highlighting the potential to obtain high-contiguity and high-quality genomes for novel species in the hospital environment using this approach (**Methods**). Rarefaction analysis of our data showed that >90% of the species and resistance gene diversity (>50% of richness) that could be sampled for sites in this study was captured by our sample size (**Suppl. Fig. S9**), while significant additional diversity remains to be captured for plasmids and HAI-associated strains (**Suppl. Note 3**). In addition, this analysis revealed that future surveys of resistance gene families in the hospital environment could be done with much fewer samples (∼50), making more regular surveys feasible, affordable and potentially actionable.

As plasmids and phages serve as an important medium for the evolution and spread of ARGs and emergence of multidrug resistant bacteria^46-49^, we identified sequences belonging to them in our genomic database and further characterized them (**Methods**). In total, we recovered 696Mbp of putative plasmid sequences (5910 closed/circular sequences and 493Mbp of linear fragments) and 63Mbp of phage sequences (1693 sequences, of which 277 are circular), with most not present in existing databases for complete plasmids^50^ or phages^51^ (>90%, 1505/1588 plasmid clusters and 501/557 phage clusters; **Methods**) despite being commonly distributed in the hospital (**Suppl. Fig. S10**), highlighting the unexplored genetic diversity in the hospital environment. Many closed plasmid sequences were >100kbp long (>9%, n=536), repeat rich and present at low abundances in the microbiome, impeding their characterization using Illumina shotgun metagenomics. In particular, we noted the presence of a group of large mecA carrying plasmids that also contained antiseptic and disinfectant resistance genes (qacA or qacC), a combination that is not represented in existing databases^50^, but is in agreement with high biocide-resistant rates for MRSA that has been reported in clinical settings^52,53^. Remarkably, one of these plasmids has genes belonging to several additional ARG classes that have not been described before in combination (e.g. dfrC, lnuA and aac6-Aph2), highlighting the value of closed plasmid sequences for detailed characterization of novel resistance gene combinations.

The availability of a large collection of highly contiguous plasmid (closed) and chromosomal (megabase-pair contigs) genome assemblies from the hospital environment allowed us to do genomic relatedness (with environmental and patient strains) and structural analysis (common gene cassettes and exchange across cassettes), as we discuss in the following sections. We first analyzed the evolutionary relationships between genomes from the hospital environment, with previously used thresholds of average nucleotide identity (ANI) to define strain-level^54^ (>99.9% ANI), derivative^16^ (>99.99% ANI) and direct-transfer^26^ (>99.999% ANI) genome clusters, and understand their spatio-temporal distribution. For many species, a diverse set of genome clusters were observed to be distributed across the hospital environment (from 6 clusters for *Pseudomonas aeruginosa* to 46 clusters for *S. epidermidis*; **Fig. 3d** and **Suppl. Fig. S11**). A subset of them were frequently detected at multiple sites and ward types in the hospital environment, and these were also significantly enriched for those detected in the 1^st^ and 2^nd^ timepoints (Fisher’s exact test p-value<1.5×10^−9^). Even at the most stringent threshold (direct-transfer), a substantial fraction of genomes observed in the 3^rd^ timepoint (1.5 years later) clustered with genomes from the first two timepoints for various species (92% for *E. anopheles* and as few as 5 SNPs; 20% for *S. marcescens* and as few as 16 SNPs; 21% for *S. haemolyticus* and as few as 8 SNPs), emphasizing the stability of environmental pathogenomes.

Overlaying antibiotic resistance information with these patterns, we noted an enrichment of multi-antibiotic resistance among strains that are widely distributed across the hospital environment through space and time (>2 antibiotics; Fisher’s exact test p-value<3×10^−8^; **Fig. 3d** and **Suppl. Fig. S12**). This was also consistently observed across several common nosocomial pathogens in the hospital environment (Fisher’s exact test p-value: 1.6×10^−2^ for *S. aureus*, 2.3×10^−3^ for *S. epidermidis*, 3.7×10^−3^ for *Enterococcus faecalis*, and 5.0×10^−2^ for *Acinetobacter baumannii*). For a subset of species (*S. aureus, S. epidermidis* and *A. baumannii*) we additionally did Illumina sequencing to generate hybrid assemblies and reliably detect derivative and direct-transfer relationships between genomes (**Methods**). Genomes that were related across the first 2 timepoints based on these stringent criteria continued to be significantly enriched for multidrug resistance (Binomial test p-value<10^−5^ for all species and both thresholds), and were enriched in the third sampling timepoint (derivative clusters, Binomial test p-value: 0.028 for *S. epidermidis* and 5.0×10^−5^ for *S. aureus*), highlighting the presence of stable, viable environmental reservoirs for common HAIs and the need to further understand the mechanisms that contribute to enrichment of multidrug resistant strains^55,56^.

### Diversity and dynamics of ARG cassettes in the hospital environment

With growing multidrug resistance, the specific combination of ARGs harbored in a strain is important to know from a clinical perspective. In the context of the hospital environment, little is known about the diversity of gene combinations and how often genes are exchanged across genomic cassettes and plasmids. Analyzing our database of 2347 high-contiguity genomes and 5910 closed plasmids together with existing genomic databases, we found that 34% of the ARG combinations observed in the hospital environment were novel (255/752) (**Suppl. File 4**). Certain ARG combinations have obvious clinical significance e.g. we noted the co-occurrence of the mecA gene (methicillin resistance) with fosB gene (fosfomycin resistance) in several environmental *S. aureus* strains, an observation that is concerning given the potential utility of fosfomycin for treating methicillin resistant *S. aureus* infections^57^. Notably, we detected Enterobacteriaceae*-*associated genes that can confer resistance to gentamycin (e.g. aac3-IIa), fosfomycin (e.g. fosA and fosA2) and colistin (e.g. mcr1), all last resort antibiotics for treating CRE infections. Two Enterobacteriaceae*-*associated plasmids, one carrying a fosA gene and the other carrying a mcr1 gene were obtained from the same bedside locker sample, highlighting the potential reservoir in the hospital environment for emergence of co-resistance to colistin and fosfomycin. Another Enterobacteriaceae*-*associated plasmid carried a rifampicin resistance gene (arr), a telling observation given the growing interest to use rifampicin in combination treatments for a variety of gram negative (e.g. *Acinetobacter baumannii*^58,59^) infections in hospitals.

We next identified common resistance gene pairs in close genomic proximity (<10kbp apart) to identify chromosomal resistance gene cassettes that may serve as the unit of evolution, co-regulation and gene exchange (**Methods**). Chromosomal cassettes for resistance genes were generally small (2-6 genes, average=3) and specific to a species, though two large cassettes carrying extended-spectrum beta-lactamases were found to overlap very well for *K. pneumonia* and *Enterobacter cloacae* (KpnC1, KpnC2 and EclC1, EclC2; KpnC3 and EclC3; **Fig. 4a** and **Suppl. File 5**). Selective pressure from the rampant use of beta-lactams and plasmid-mediated transmission is likely to have contributed to the sharing of these large cassettes across species. In general, cassettes for gram-negative species were larger and more stable (solid lines to genes) while those for gram-positives were smaller with many variably present members (dashed lines to genes). The largest shared cassette among gram-positives (aminoglycoside-streptothricin resistance; ant6-Ia, sat4A and aph3-III) was seen in *Enterococcus* and *Staphylococcus* species, but with no discernible signals of mobile elements^60^. While most genes were observed to be stably present in cassettes (solid lines; except for a few e.g. tetK, far1, catA), exchange of genes across cassettes were rarely observed (e.g. blaZ), indicating that chromosomal cassettes tend to be relatively fixed.

**Figure 4:**
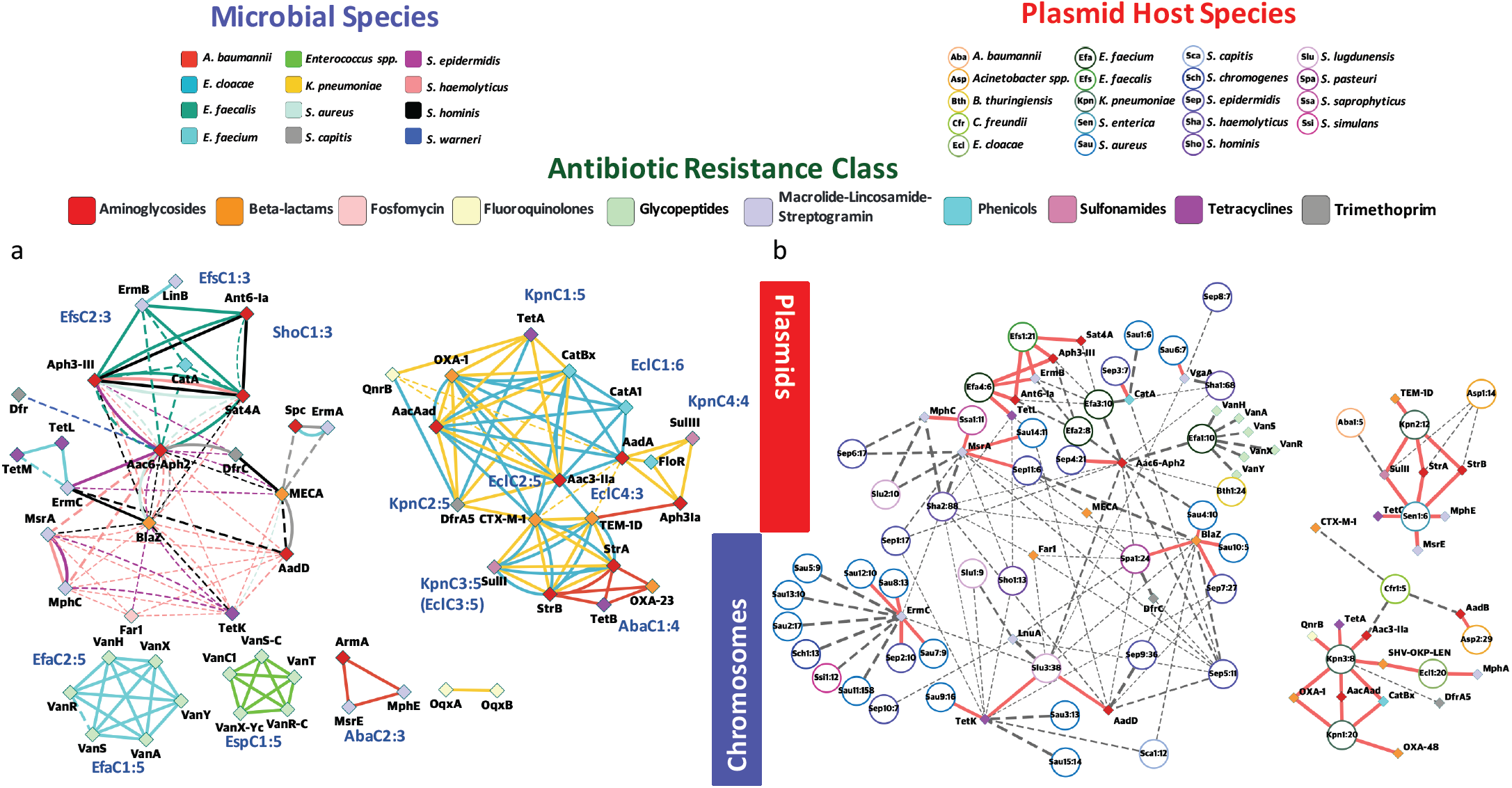
Species distribution and genomic proximity of drug resistance genes in the hospital environment microbiome. Genomic proximity network and clustering of ARGs based on 2347 microbial genomes and 5910 closed plasmids obtained from the hospital environment. (a) Multigraph of genomic proximity between ARGs, where edges indicate gene pairs found <10kbp apart in the genomes for a species (not including plasmids). Edges are colored according to species and line widths indicate frequency of occurrence of gene pairs (normalized by count for rarer gene) with frequencies >80% marked with solid lines. Solid line cliques in each species were used to define cassettes and assign names (**Suppl. File 5**) and the number after the colon sign indicates clique size. Genes are colored according to their respective antibiotic classes. (b) Circles represent different plasmid clusters (95% identity) and corresponding ARGs belonging to them are connected by edges and indicated by diamonds. Plasmid nodes are labelled based on a three letter short form for the host species and a number (e.g. Kpn1 for a *K. pneumoniae* plasmid) and the number after the colon sign indicates the number of representative of the plasmid family that were observed in the database. Edges are weighted by the frequency at which a gene is present in a plasmid and frequencies >80% are indicated with red solid lines. Genes and backbones are color-coded according to their respective ARG classes and inferred host species for ease of reference.

Performing similar analysis for the closed plasmid sequences identified in this study, we first clustered them into shared plasmid backbones and annotated them for their known hosts (identity ≥95%; **Methods**). Analyzing ARGs in the context of these backbones, we found that many resistance genes are variably present in backbones (93/143 genes), and for genes that are stably found in one backbone, a high frequency are also variably present in another backbone (19/31 genes), highlighting the dynamic nature of resistance genes combinations in the plasmids that we recovered in the hospital environment (**Fig. 4b**). In this background, we observed a few gene combinations that were stably present in multiple plasmid backbones, pointing to strong selection for co-existence. For example, the genes strA, strB (streptomycin resistance) and sulII (sulfonamide resistance) co-occur in two distinct backbones (Sen1 and Kpn2, sequence overlap <54%) as a signature from past co-administration of streptomycin and sulphonamides for clinical use^61,62^. Similarly, while aminoglycoside resistance genes such as aadD and aac6-Aph2 are widely and variably distributed across plasmid backbones, the genes ant6-Ia and aph3-III are stably shared by two distinct backbones (Efa4 and Efs1, sequence overlap <11%) indicating that they may provide synergistic resistance to aminoglycosides by catalysing different modifications. Interestingly, we noted that genes that are widely distributed across plasmids (e.g. tetK, far1 and blaZ) can come together in a novel, clinically relevant plasmid backbone (**Fig. 4b**; Slu3, with 38 sequences in our database) as described for a cytotoxin producing MRSA strain^63^. While the previously isolated strain was resistant to fusidic acid and tetracycline, but susceptible to erythromycin and clindamycin, we noted the presence of a common plasmid backbone in our database (Sha2 with 88 sequences) that carries a novel combination of resistance genes for all 4 antibiotics (**Fig. 4b** and **Suppl. File 4**). Similarly, we found that resistance genes found in phages, such as aac6-Aph2 and far1, tend to be more widely present (**Fig. 4** and **Suppl. Fig. S13a**), with evidence for recent phage-mediated dissemination of far1 across *Staphylococcus* species (**Suppl. Fig. S13b**). In general, we observed that ARGs found in plasmids tend to have more genes in close proximity (<10kbp apart) in chromosomes than chromosome-exclusive genes (Wilcoxon test p-value=6×10^−7^), characteristic of higher gene mobility and shuffling for plasmid-associated genes. Thus the plasmid backbones seen in the hospital environment likely represent a more plastic framework to generate diverse ARG combinations, many of which are not seen in genomic cassettes (25%), despite the strong overlap in the complement of resistance genes that they harbour (84% of plasmid genes).

### Hospital environment strains overlapping with patient isolates are globally disseminated and enriched for multidrug resistance

The availability of a large database of genomes from many species in the hospital environment, an obvious hub for patient colonization, prompted us to ask the question: “How are environmental strains related to patient colonizing strains?”. To examine this relationship, we jointly analyzed environmental strains and patient isolates for different species to construct phylogenetic trees (**Fig. 5**). We started by analyzing Singaporean *E. anophelis* isolates from a 2012 HAI outbreak^64^ (n=10) and an additional set of patients isolates collected from 2009-2012 (n=52; **Fig. 5a**; **Methods)**. Strikingly, despite sampling from different hospitals in Singapore and after a span of 5-8 years, patient associated genomes matched environmental genomes with as few as 16 SNPs (s1; average nucleotide identity, ANI 99.9996%). The environmental *E. anophelis* genomes in our studies primarily originated from hospital sinks, which as we noted earlier tends to have a stable community, indicating that these strains may have originated from a common reservoir upstream of water piping systems^54^. In addition, we noted that *E. anophelis* clusters shared between patients and the environment were also detected at the 3^rd^ timepoint sampling 1.5 years later (ANI >99.999%, corresponding to direct-transfer), and exhibited resistance to more antibiotics (fold-change=1.25×, one-sided Wilcoxon p-value=0.059).

**Figure 5:**
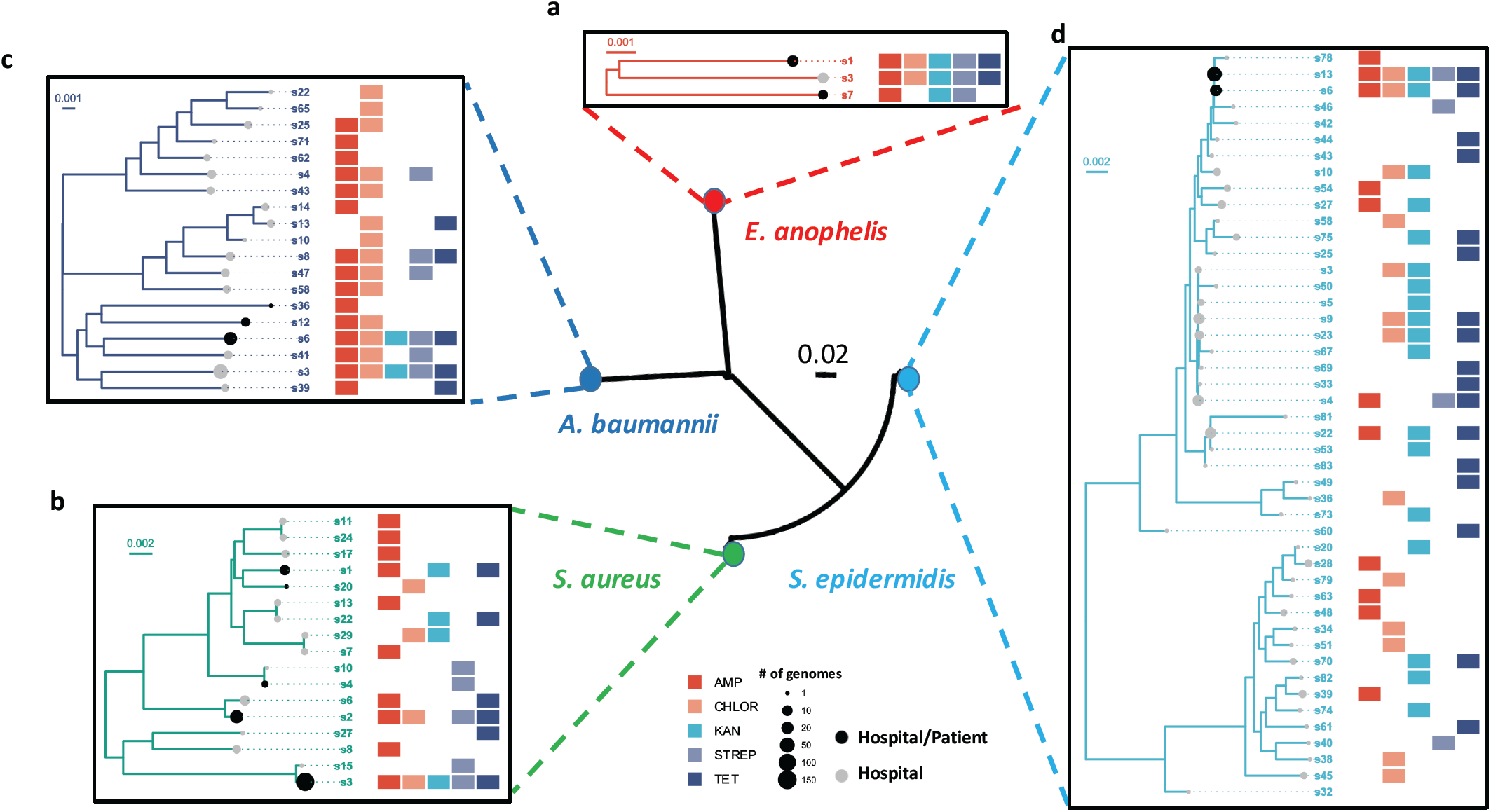
Multi-species analysis of phylogenetic relationships between environmental and patient genomes. Phylogenies depicting the evolutionary relationships between derivative clusters (>99.99% ANI), with each node representing the consensus genome for a cluster, for (a) *Elizabethkingia anophelis* from a nosocomial outbreak in Singapore in 2012 and other patient isolates, (b) patient colonizing *Staphylococcus aureus* from a 2009-2011 surveillance study in Singaporean hospitals, (c) infectious *Acinetobacter baumannii* isolates from patients in two major Kuwaiti hospitals and Singaporean patient isolates, and (d) recent globally disseminated multi-drug resistant *Staphylococcus epidermidis* lineages, together with environmental genomes for corresponding species from this study. While subfigures a and b highlight the close relationships between strains circulating in Singaporean hospitals that are up to 8 years apart, subfigures c and d reveal the global dissemination of several lineages. Note that the matrices next to the trees indicate the antibiotic resistance profiles for corresponding derivative clusters. Scale bars depict the number of substitutions per site in the core alignment. For all species tested, clusters shared between environmental and patient genomes were enriched for multidrug resistance.

Encouraged by the high-similarity match observed for *E. anophelis*, we analyzed *S. aureus* genomes (n=221) obtained from a surveillance study of hospitalized patients in the same facility almost a decade ago^65^. These strains were found to match 5 out of 17 strains that were obtained in the current study, with genomes having as few as 39 SNPs and average nucleotide identity as high as 99.9985% (**Fig. 5b**). The co-occurrence of patient and environmental genomes was found to be significantly enriched in multidrug resistant clusters at the derivative genomes threshold (e.g. s1, s2 and s3; Binomial test p-value<10^−15^). These clusters were also enriched for genomes detected in the 3^rd^ timepoint sampling (Binomial test p-value<2×10^−7^) with <60 SNPs (ANI 99.998%) distinguishing them from genomes in the 1^st^ and 2^nd^ timepoints, highlighting the stability of antibiotic resistant derivative clusters in the hospital environment.

To further extend our observations, we sequenced *A. baumannii* patient isolates (n=108) from a hospital surveillance cohort in Singapore that was established >8 years ago. Patient isolate genomes from this cohort were found to have high identity to some of our environmental genomes (99.995% ANI, cluster s6), while being temporally separated by almost a decade. In addition, patient isolates that overlapped with environmental genomes were enriched for multidrug resistance status (derivative clusters; Binomial test p-value<4×10^−3^). Extending our analysis to a regional context, we analyzed genomes (n=36) for *A. baumannii* patient isolates from two major Kuwaiti hospitals^66^ along with our environmental genomes (**Fig. 5c**), and found that a derivative cluster shared between environmental and Singaporean patient genomes exhibited resistance to all 5 antibiotics in this study and also included Kuwaiti isolates at high identity (s6, ANI >99.99%). Our data thus highlights the presence of multidrug resistant *A. baumannii* derivative clusters in hospital environments that are persistent, enriched in the overlap with patient isolates and globally disseminated.

Similar patterns were observed recently for *S. epidermidis* lineages (ST2/ST2-mixed) which seem to have disseminated across the globe within a short period (n=229)^40^. We confirmed the detection of these rifampicin resistant^40^ lineages in our data (**Fig. 5d**) with as few as 80 SNPs (ANI 99.997%) to our hospital environment genomes. In addition, one other lineage (ST16) not known to be globally disseminated (isolated from a patient sample in USA^40^) was also represented by a genome in our database that is similar at the derivative threshold (ANI 99.991%). Finally, we again found that the overlap between *S. epidermidis* patient isolates (surveillance samples from Austin Health in Australia^40^) and environmental genomes from this study was enriched for multidrug resistance in derivative clusters (Binomial test p-value<4×10^−12^). Together with the observation that multi-antibiotic resistant strains are persistent and widely distributed across the hospital (**Fig. 3d** and **Suppl. Fig. 11**), this data points to selective advantages for multi-drug resistant organisms to persist and spread in hospital environments and patients.

## Discussion

While the importance of a well-designed built-environment for preventing infections in hospitals is well appreciated^67^, the use of shotgun metagenomic approaches for surveying microbial communities established in functioning hospitals or medical environments remains underexplored^18^. A large baseline survey such as the one conducted here can help provide a reference map (potentially visualized using 3D tools, e.g. https://github.com/csb5/hospital_microbiome_explorer) that can then be updated based on periodic scans whose frequency and locations can be informed by the initial survey. For example, the turnover score and specificity of a site can help determine whether and how frequently it should be sampled. Variations in human influence scores could also help fine-tune cleaning practices, and the distribution of specific opportunistic pathogens of concern could be informative in an outbreak setting for infection control. As genomics-guided infection control is further refined, the knowledge gained could then feed back into better hospital designs. With further improvements in cost and ease of short-read sequencing, we anticipate that hospital-wide surveys will be increasingly feasible, will provide valuable information for infection control, and eventually be part of routine practice.

The two microbial community types observed in this study, highlight the distinct niches found in hospital environments compared to other urban environments, and provide an organizing principle for further study. For example, while many pathogens are substantially enriched in the hospital environment, this appears to be more a feature of CTA sites which also have a greater diversity of ARGs (**Suppl. Fig. S7c, d**). From a clinical perspective, recent infection control studies have primarily focused on sites associated with wash areas (sinks, showers etc.; CTB sites), as in many cases the presumptive pathogens for an outbreak have been isolated from such sites^54^. This focus on community type B sites is in line with them being composed of biofilm-forming bacteria and having the ability to harbor viable reservoirs for extended periods of time (e.g. in the plumbing system). Our data however suggests that many medically-relevant species (e.g. *K. pneumoniae*), ARGs (e.g. carbapenemases such as oxa-23) and resistance plasmids (present in >85% of CTA sites) are more frequently harbored in community type A sites. While CTA sites have higher microbiome turnover, the ability to detect very similar strains over long durations of time indicates that they may also have distinct reservoirs (e.g. in ventilation or air-conditioning ducts), and that culture-based screening protocols may bias against sites that have lower biomass or are variably colonized. Combining the strengths of metagenomic sequencing and culturing may therefore be needed to more systematically explore the source of outbreaks.

Large-scale genomic studies of nosocomial pathogens through strain isolation can be a laborious and time-consuming process, while as shown here, direct shotgun metagenomics will often not provide detailed genetic information for species present at low relative abundances. The intermediate approach proposed here attempts to address both issues. The use of culture-based enrichment allows us to effectively shift the distribution away from abundant species such as *C. acnes*, and towards nosocomial pathogens that are typically at low relative abundances (e.g. *K. pneumoniae, S. aureus* and *A. baumannii*), while also allowing for functional selection such as for antibiotic resistance as we did here. Its combination with long-read metagenomic sequencing is then powerful as it allows us to directly recover high-contiguity genomes (chromosomal, plasmid and phage) without a tedious isolation step. With further automation (e.g. in library preparation) this workflow is suitable for high-throughput analysis and amenable for wider surveillance, in line with the vision to achieve precision epidemiology for infectious diseases^68^. Future improvements in nanopore sequencing throughput, the ability to work with lower DNA input, and use in remote settings, could help accelerate time-to-answer by reducing the culturing period, or eliminating it altogether.

The availability of a database of high-contiguity assemblies, with >8Gbp of sequence, 2347 genomes, 1693 phage sequences and 5910 closed plasmids, provides a unique resource for studying transmission patterns of strains and diversity of resistance gene cassettes across species in the hospital environment. Based on this resource, we observed that multidrug resistant strains are preferentially distributed and persistent in hospital environments across a range of species. This represents a worrisome pattern, with several possible explanations that deserve further investigation. One plausible scenario is that hospital environments are repeatedly seeded by multidrug resistant strains that are preferentially persistent in the community (humans or environment). This scenario seems less likely to be generally true given that some of the species where we observe this pattern are rarely found in humans (e.g. *E. anophelis, A. baumannii*), and based on the observations here that other urban environmental microbiomes are distinct from hospital environments in terms of composition, frequency at which they harbor species associated with nosocomial infections, and diversity and abundance of ARGs. Nevertheless, the data in this study does not rule out the possibility that urban environments, (i) harbor pathogens and resistant strains at lower abundances compared to hospitals, and (ii) that antibiotic resistant strains are also widespread and persistent in these environments. Another potential hypothesis is that hospital cleaning measures may be selecting for more antibiotic-resistant organisms^69,70^, a model that is also supported by the presence of multiple copies of disinfectant resistance genes in the widely distributed, multidrug resistant *S. aureus* strains in our study. Comparisons with surveys from built-environments that are intensively cleaned but do not house patients (e.g. operating rooms) or are not intensively cleaned but have high patient traffic (e.g. waiting areas in clinics) might help explore this model further. Studies across wards or hospitals with different cleaning protocols could also be illuminating for understanding how ARG reservoirs in hospitals can be shaped by infection control practices^71,72^.

Despite its importance as the epicenter for the battle against growing antibiotic resistance^1^, hospital environments have received relatively little attention compared to studies on agricultural and animal farms^18^. Our analysis highlights that hospital environments harbor significant uncharacterized genetic diversity in terms of microbial species (13 novel species) and ARG combinations (255). This reservoir can be the origin of new opportunistic infections and serve as fertile ground for evolution of ARG combinations that are of clinical concern (e.g. colistin and fosfomycin resistance). This can be further facilitated by the presence of resistance gene carrying plasmids that serve as a vehicle for gene transfer across species^73^, and were commonly found in the hospital environment (n=1400). The development and use of anti-plasmid agents^74,75^ could thus be a complimentary strategy to reduce the spread of ARGs through hospital environments.

While other studies have focused on patient isolates^76^, the genetic relatedness between environmental and patient-colonizing strains in hospitals is important to study for understanding the potential risk that environmental strains carry for causing infections^24,26^. For strains that are contemporary and co-located, high genetic relatedness between a subset of environmental and patient strains is expected. The observation that highly similar genomes can be obtained despite being temporally separated by >8 years suggests that large reservoirs of nosocomial multidrug resistant strains are being maintained with limited diversification. The identification and elimination of these reservoirs may help reduce the incidence of corresponding infections and the risk from maintenance of antibiotic resistance properties. Another distinct observation is the high genetic similarity between MDRO strains observed in Singaporean hospitals and those obtained from patients around the world. The consistency of these patterns across species emphasizes the global dissemination of newly emerged MDRO lineages and the role of hospital environments in this deserves investigation, in conjunction with global environmental metagenomic datasets^37^.

Taken together, our data points to selective advantages for multidrug resistant organisms to persist and spread in hospital environments (**Fig. 3d** and **Suppl. Fig. S11**) and be shared with patients (**Fig. 5**). The *S. aureus* derivative clusters that persisted in the hospital environment across time are enriched in virulence factors (1.5×, one-sided Wilcoxon p-value=0.015) and have as many as three copies of disinfectant resistance genes^52,53^ (**Fig. 3d**). This may enable them to successfully colonize both hospital environments and patients, and thus be readily transferred between them. This points to a vicious cycle where disinfectant resistance, antibiotic resistance and virulence may in turn be selected for, enriching for strains that are adept at colonizing both niches in the presence of depleted microbial competition, and offers an explanation for the high incidence of multidrug resistant HAIs worldwide despite increased surveillance and aggressive cleaning measures in hospitals^77-80^.

## Methods

### Sample collection and storage

Environmental swabs were collected from Tan Tock Seng Hospital (TTSH), a major tertiary care hospital with >2000 patient visits every day that serves as the national referral center for communicable diseases in Singapore. Sampling was carried out in November 2017 and in May 2019. Samples for the first timepoint being collected in 2 days and the second timepoint in 3 days, with 1 week separating the timepoints. The third timepoint was 1.5 years later, collected in 4 days across 2 weeks. Samples were collected from isolation rooms (1 bed, typically warding CRE colonized patients), MDRO wards (5 beds, typically warding MRSA colonized patients) and standard wards (5 beds), at 7 different sites, including aerator, sink trap, bed rail, bedside locker, cardiac table, pulse oxymeter and door handle (**Fig. 1** and **Suppl. File 1**). Standard cleaning protocols at TTSH require that high-touch areas and sinks are cleaned daily (chlorine 5000ppm and cleaning detergent, respectively; excluding beds that are cleaned upon patient discharge). Isohelix DNA Buccal Swabs (SK-4S) were used for sampling carried out based on MetaSUB protocols^37^. Briefly, a total of 4 swabs were collected with 1 swab (for culturing) moistened with 1X phosphate buffer saline (PBS, pH 7.2) and 3 swabs (2 for metagenomic DNA isolation, 1 for storage) moistened with DNA/RNA shield (Zymo Research, Cat. No. ZYR.R1100-250). Swabbing was performed for 2 min in each site and stored in respective storage liquids (i.e. 1X PBS, pH 7.2 or Zymo DNA/RNA shield). Swabs in PBS were placed on ice and sent for culturing while the other swabs were transported back at room temperature to the laboratory and stored at -80°C. In total, 1752 swabs were collected from 179 sites in the hospital at 3 timepoints, representing 438 unique samples. Swabs were also collected from an office environment (Genome Institute of Singapore) with sites selected to approximately match those collected in the hospital (aerator, sink trap, chair handle, office desk, keyboard and door handle; n=30; **Suppl. File 1**). MetaSUB Singapore samples were collected from high-touch surfaces in different parts of the city and analyzed based on MetaSUB protocols as described in Danko et al^37^ (n=99; **Suppl. File 1**).

### DNA extraction from swabs

DNA was extracted from swabs using a bead-beating and automated DNA purification system. Briefly, 300 µL of lysis buffer was added to lysing matrix E tubes (MP Biomedicals, Cat. No. 116914500). Samples were homogenized using the FastPrep-24 instrument at 6 m/s for 40s prior to centrifugation at maximum speed for 5 min. The supernatant was treated with Proteinase K (Qiagen Singapore Pte. Ltd, Cat. No. 19133) for 20 min at 56°C before DNA was purified with Maxwell® RSC Blood DNA Kit (Promega Pte. Ltd., Cat. No. AS1400). DNA concentration was quantified using Qubit® 2.0 fluorometer, prepared with Qubit® dsDNA HS Assay Kit (Life Technologies Holdings Pte. Ltd., Cat. No. Q32854). DNA extraction from backup swabs was carried out for samples with insufficient amounts of DNA. Samples that still had less than 0.5ng of DNA were excluded for library preparation (10/438).

### Illumina library preparation

Extracted DNA was sheared using Adaptive Focused Acoustics™ (Covaris) with the following parameters: 240s, Duty Factor: 30, PIP: 450, 200 cycles/burst. Metagenomic libraries for the first two timepoints were prepared with NEBNext® Ultra™ DNA Kit (New England Biolabs, Cat. No. E7370) according to manufacturer’s instructions. Paired-end sequencing (2×101bp reads) was performed on the Illumina HiSeq2500 platform. For the third timepoint, metagenomic libraries were prepared using NEBNext® Ultra™ II DNA Library Prep Kit (New England Biolabs, Cat. No. E7645) according to manufacturer’s instructions. Paired-end sequencing (2×151bp reads) was performed on the Illumina HiSeq4K platform.

### Culture enrichment

Following MetaSUB protocols, swabs were directly incubated with 7 mL of Brain Heart Infusion (BHI) broth (Thermo Scientific Microbiology, Cat. No. CM1135B) at 37°C till turbidity was observed (14-16 h for >95% of samples) or up to a maximum of 48 h. Culture tubes were centrifuged at 3200 g for 12 min. For the first two time points, cell pellets were re-suspended with 550 µL of 1× PBS while the cell pellets for the third time point were re-suspended with 1 mL of 1× PBS. Fifty microliters of re-suspended cultures were then plated on each of the 6 agar plates (BHI without antibiotics, Ampicillin 100 µg/mL - AMP, Streptomycin sulfate 100 µg/mL - STREP, Tetracycline 10 µg/mL - TET, Kanamycin 50 µg/mL – KAN and Chloramphenicol 35 µg/mL - CHLOR) and incubated overnight at 37°C. Cells were harvested by a plate sweep and were pelleted down by centrifugation at 8000 *g* for 15 min at 4 °C for the first two time points. For the third time point, a loopful of harvested cells was streaked out on an antibiotic-free BHI plate to obtain single colonies for whole genome sequencing. Plates were only excluded if no cells were growing on the plates or when the growth was insufficient to generate enough DNA for sequencing.

### DNA extraction from enrichment cultures

Frozen cells were thawed on ice and manually mixed with a wide bore tip. A volume of 30-50 µL of cells was re-suspended in 100 µL of 1× PBS, pH 7.4. Twenty microliters of suspended cells were added to 20 µL of metapolyzyme (6.7 µg/µL) (Sigma Aldrich, Cat. No. MAC4L). The mixture was incubated at 35°C for 4 h. RNase treatment was carried out by adding 350 µL of 1× TE buffer and 10 µL RNase A (4 mg/µL) and incubated on a rotator for 10 min at room temperature. DNA was extracted with Maxwell® RSC cultured cells kit (Promega Pte. Ltd., Cat. No. AS1620). DNA was cleaned up and concentrated with 0.4× Agencourt AMPure XP beads (Beckman Coulter, Cat. No. A63882). DNA purity and concentration was measured with Nanodrop and Qubit® fluorometer. DNA integrity was assessed on a 0.5% agarose gel. DNA samples with the following quality measurements were selected for nanopore sequencing (DNA amount >400 ng, A260/280: 1.8-2.0, A260/230: 1.7-3.0, Qubit:Nanodrop: 0.7-1.3, DNA integrity on 0.5% agarose gel: >1 kb). The Qubit:Nanodrop ratio was used to estimate and control the amount of single-stranded DNA in the sample, and ensure successful nanopore sequencing.

### Collection and testing of bacterial isolates from patients

*Elizabethkingia anophelis* isolates (n=52) were obtained from consecutive positive blood cultures and respiratory samples collected in a 3-year period (2009-2012) at the National University Hospital in Singapore (DSRB reference number 2017/00879). *Acinetobacter baumannii* complex isolates (n=108) were consecutively obtained from all clinical specimens (including blood, tissue, respiratory and urine samples) sent for routine bacterial culture between February 2009 and May 2009 at the Singapore General Hospital Diagnostic Bacteriology Laboratory (de-identified and archived and hence no IRB approval was required). Antibiotic susceptibility testing for *E. anopheles* isolates was done with 13 antimicrobial agents (cefotaxime, ceftazadime, cefepime, imipenem, meropenem, ampicillin-sulbactam, piperacillin/tazobactam, tigecycline, gentamicin, nalidixic acid, ciprofloxacin, levofloxacin and trimethoprim/sulfamethoxazole) using Etest strips (bioMérieux). Minimum inhibitory concentrations (MICs) were interpreted according to the Clinical and Laboratory Standards Institute (CLSI) guidelines for non-Enterobacteriaceae Gram-negative bacilli (Performance standards for antimicrobial susceptibility testing, M100-S22, CLSI 2012; **Suppl. File 6**). Antibiotics that all strains were resistant to were excluded from statistical analysis. Antibiotic susceptibility testing for *A. baumannii* complex isolates was done with 11 antimicrobial agents (ampicillin/sulbactam, piperacillin/tazobactam, cefepime, imipenem, gentamicin, amikacin, ciprofloxacin, levofloxacin, trimethoprim/sulfamethoxazole, minocycline and polymixin B). Polymixin B susceptibility testing was performed using Etest strips (bioMérieux) and disk diffusion was performed for all other antimicrobial agents. Polymixin B minimum inhibitory concentrations (MICs) and zone diameters for all other tested agents were interpreted in accordance to Clinical and Laboratory Standards Institute (CLSI) breakpoints for *Acinetobacter* spp. (Performance standards for antimicrobial susceptibility testing, M100-S19, CLSI 2009; **Suppl. File 6**). Multidrug resistant status for patient isolates was defined according to CDC guidelines (https://www.cdc.gov/nhsn/pdfs/ps-analysis-resources/phenotype_definitions.pdf).

### DNA extraction for bacterial isolates

Cell pellets were allowed to thaw slowly on ice and re-suspended in 400 µL of ATL buffer (Qiagen Singapore Pte. Ltd, Cat. No. 19076). Cells were lysed in Lysing Matrix E tubes (MP Biomedicals, Cat. No. SKU 116914500) on a vortex adapter at maximum speed for 10 min. Cell lysates were centrifuged at 16,000 *g* for 5 min and supernatant was treated with 4 µL of RNase A (100 mg/mL) (Qiagen Singapore Pte. Ltd, Cat. No. 19101), gently mixed by flicking of tube and incubated at room temperature for 2 min. The cell lysate was further treated with 25 µL of proteinase K (20 mg/mL) (Qiagen Singapore Pte. Ltd, Cat. No. 19133), gently mixed by flicking of tube and incubated at 56 °C for 20 min. DNA was purified twice using 1× volume of AMPure XP beads (Beckman Coulter, Cat. No. A63882) with slight modifications to the manufacturer’s protocol. All mixing steps were replaced with gentle flicking of tube and incubation on the hula rotor for gentle mixing. Fresh 70% ethanol was prepared for washing and magnetic beads were incubated on a 37 °C heat block for 3–5 min to dry off residual ethanol. The quality and quantity of DNA was assessed using Nanodrop, Qubit® fluorometer and 0.5% agarose gel. Samples that were unable to pass the following criteria were omitted from sequencing: DNA amount measured by qubit >510 ng, DNA concentration measured by qubit >11 ng/µL, A260/280 ratio between 1.7-2.0, A260/230 ratio between 1.5-3.3 and DNA length >1 kb. Purified DNA was stored at 4 °C.

### Nanopore library preparation

DNA was prepared with either 1D^2^ sequencing kit (SQK-LSK308) or 1D sequencing kit (SQK-LSK108 or SQK-LSK109) together with native barcoding kit (EXP-NBD103 or EXP-NBD104 and EXP-NBD114) according to the native barcoding genomic DNA protocol. DNA was not sheared and was used directly for DNA repair and end-prep. Both native barcode ligation and adapter ligation steps were extended to 30 min instead of 10 min. In addition, to maximize library yields, more than 700 ng of pooled sample (where possible) was used for adapter ligation. Samples were multiplexed (9-12 samples per pool for culture enriched samples and 24 samples per pool for isolates) and sequenced with either MIN106, MIN106D or MIN107 flowcells on a GridION machine.

### Taxonomic and resistome profiling with Illumina shotgun metagenomic data

Illumina shotgun metagenomic sequencing reads were processed using a Snakemake^81^ pipeline (https://github.com/gis-rpd/pipelines/tree/master/metagenomics/shotgun-metagenomics). Briefly, raw reads were filtered to remove low quality bases using skewer^82^ (v0.2.2; -q 3 -l 30 -n) and human reads were removed by mapping to the hg19 reference using BWA-MEM^83^ (v0.7.10-r789). The remaining microbial reads were profiled with MetaPhlAn2^84^ (v2.6.0) and SRST2^85^ (v0.1.4; --min_coverage 100, hits with identity <99% were filtered out) for taxonomic and ARG abundances, respectively. Microbial reads were also assembled using MEGAHIT^84^ (v1.0.4-beta; default parameters) for comparison to nanopore assemblies. Site specificity score was computed as the z-score for the closest taxonomic profile for a sample (Bray-Curtis dissimilarity) among physically proximal sites (in the same room/cubicle and same timepoint), compared to the distribution of Bray-Curtis dissimilarities across all samples of a site (e.g. all bed rails). Results based on analysis of taxonomic and resistome profiles were obtained for each timepoint independently and compared across timepoints to check for consistency and filter out potential sequencing artefacts^34^.

### Removal of likely contaminant species

Likely contaminant species were identified based on batch and correlation analysis^*34*^ (**Suppl. Note 2**) and removed from species-level abundance profiles. For genus-level profiles, relative abundances of the filtered species were subtracted from the abundance of the corresponding genera for each sample. Filtered profiles were then renormalized to sum to 100% and used for all downstream analyses.

### Preprocessing of nanopore sequencing data

Raw nanopore reads were basecalled with the latest version of the basecaller available at the point of sequencing (Guppy v0.5.1 to v3.0.6 or Albacore v2.3.1 to v2.3.3 for libraries that failed live basecalling). Basecalled nanopore reads were demultiplexed and filtered for adapters with Porechop (v0.2.3; https://github.com/rrwick/Porechop) or qcat (v.1.1.0 https://github.com/nanoporetech/qcat). Sequencing statistics were summarized using SeqKit^86^ (v0.10.1). Reads were taxonomically classified with Kraken^87^ (v0.10.5-beta) against the miniKraken database (minikraken_201711_01_8GB_dustmasked) to assess the diversity of cultures on the plates.

### Genome assembly and species assignment

Nanopore reads for each plate were assembled using Canu^88^ (v1.3/v1.7; genomeSize=8m). For samples where both Illumina and Nanopore reads were available, a higher quality hybrid assembly was obtained using OPERA-MS^28^ (v0.8.3; --polish --no-gap-filling --short-read-assembler spades). Assembled contigs were mapped to the NCBI nt database with BLAST (v2.2.28), to identify microbial species or plasmid assignments according to the best BLAST hit (highest total reference coverage). Circular sequences were identified using MUMmer^89^ (v3.23; -maxmatch -nosimplify, alignments <1kbp long or with identity <95% were filtered out) as recommended in the documentation for the Canu assembler (https://canu.readthedocs.io/en/latest/faq.html#my-circular-element-is-duplicated-has-overlap). Contigs assigned to the same species were binned into genomic bins. Metagenomic Illumina reads were used to polish Canu assemblies where feasible using Pilon^90^ (v1.22, --fix indel). We noted that annotation errors were significantly reduced after polishing, and that genomic bins whose length was within 10% of the expected length met the criteria for high-quality genomes (completeness >90% and contamination <5% using CheckM^91^; v1.0.7 --reduced_tree). Genomic bins that met these criteria were therefore designated as high-quality and incomplete bins (<50% of expected length) were removed from further analysis. Genomes corresponding to novel species were identified as those with identity <95% or coverage <80% compared to known genomes (BLAST with nt) and three recent catalogs that include environmental and human microbiome assembled genomes^43-45^ (with Mash^92^). The genomes were hierarchically clustered (single-linkage with Mash distance^92^) to identify species-level clusters at 95% identity and genus level taxonomic classification was obtained using sourmash^93^. Similarly, novel circular plasmids were identified by comparing to the PLSDB^50^ database with Mash distance and identifying clusters at 99% identity (single-linkage) with no known sequence.

### Analysis of ARG combinations

ARGs were annotated to contigs by mapping them to the ARG-ANNOT^94^ database provided in SRST2 (v3) with BLAST (best hit with >90% identity and >90% reference coverage). ARG combinations present in chromosomes and plasmid sequences were considered novel when they were not found in the reference databases (nt or PLSDB^50^). Assembled circular plasmids were clustered and annotated based on their best BLAST hit with identity >95% and >60% query coverage. A bipartite graph was constructed by connecting each plasmid cluster to ARGs found in it, with edge weights representing the frequency of occurrence (clusters with <5 representatives were excluded). For each species, an ARG co-occurrence graph was created for ARGs found in the assembled genomes by connecting the ARG pairs that were found within 10kbp on the same contig (discarding ARG pairs occurring less than 5 times). Each edge was weighted by the frequency of ARG pairs divided by the minimal frequency of the two ARGs. All ARG co-occurrence graphs were merged into a final co-occurrence multigraph. The graphs were visualized using Cytoscape (v3.7.1)^95^.

### Analysis of virulence factor and biocide resistance genes

Nanopore assemblies were aligned to virulence factors in the PATRIC database^96^ (2018/12/20) with DIAMOND (v0.9.24.125; blastx --long-reads) and alignments with E-value >0.001 were filtered out. To identify biocide resistance genes, the assemblies were aligned to nucleotide sequences for the genes qacA (NC_014369.1) and qacC (NC_013339.1) with BLAST (>90% identity and >90% reference coverage).

### Analysis of phages and prophages

Phage-like-elements (phage and prophages) were identified using VirSorter^97^ (v1.0.5, phages and prophages in category 3 or with length <10kbp were filtered out). The assembled phages and reference phages from the MVP database^51^ were hierarchically clustered (single-linkage with Mash distance^92^) to identify phage clusters at 95% identity. Clusters without any phages from the reference database were considered novel. For each cluster, sub-clusters were defined at 99.9% ANI by single-linkage clustering with nucleotide identities from nucmer (--maxmatch –nosimplify, followed by dnadiff and minimum sequence overlap of 80%). Phage-like-elements were annotated using RAST^98^ (virus domain, fix frame-shifts parameters).

### Analysis of patient isolates and strain relationships

Raw reads corresponding to genomes for outbreak isolates^40,64-66^ were downloaded and assembled using the Velvet assembler^99^ (v1.2.10) with parameters optimized by Velvet Optimiser (k-mer length ranging from 81 to 127), scaffolded with OPERA-LG^100^ (v1.4.1), and gapfilled with FinIS^101^ (v0.3). Outbreak genomes from the same species were jointly analyzed with high-quality genomes from the hospital environment. To identify high confidence SNPs we adapted the method from Brooks et al^26^. Specifically, we performed pairwise alignments between genomes using nucmer, and considered genome pairs with alignment coverage >80% for ANI computation. SNPs between genome pairs were called using MUMmer’s ‘show-snps’ function and regions containing more than 1 SNP within 20bp were filtered out to mask potential artefacts from horizontal gene transfer, recombination or repeats. Finally, the genomic distance matrix (based on ANI = #SNPs/alignment size) was clustered hierarchically (single linkage) and clusters obtained at 99.99% identity for species with hybrid (Nanopore and Illumina) assemblies (or 99.9% for species with Nanopore-only assemblies). Single-linkage clustering was used to avoid having highly similar genomes being assigned to separate clusters and we confirmed that despite this, most members (99%) had an average distance to other members of the cluster below the clustering thresholds used. Antibiotic resistance profiles and multidrug resistance status (>2 antibiotic types) for each cluster were derived from the union of resistance profiles for each genome obtained in various selection plates.

For phylogenetic analysis, a consensus genome was derived for each cluster based on reference-guided alignment with nucmer (*S. aureus*: NC_020529, *S. epidermidis*: NC_004461, *E. anophelleis*: NZ_CP007547, *E. faecalis*: NC_017312, *E. faecium*: NC_017960, *P. aeruginosa*: NC_018080, *K. pneumoniae*: NC_018522, *A. baumannii*: NC_009085) and the cons utility in the EMBOSS suite^102^. Maximum likelihood phylogenetic trees were constructed for each species with Parsnp^103^ (v1.2; -c –x; accounting for recombination events using PhiPack^104^) based on consensus genomes for each cluster, where multiple sequence alignments for each species varied in length from 0.6Mbp (*S. epidermidis*) to 5.1Mbp (*P. aeruginosa*). For the species-level tree, full-length 16S rRNA sequences (*S. epidermidis*: L37605.1, *S. aureus*: NR_118997.2, *E. anophelis*: NR_116021.1 and *A. baumannii*: NR_026206.1) were aligned with MAFFT^106^ (v7.154b, default parameters) and the phylogeny was determined using FastTree2^107^ (v2.1.8, default parameters). The trees were visualized using the ‘ggtree’ R package^105^. Strain distributions across sites were visualized with the ‘HiveR’ R package (academic.depauw.edu/∼hanson/HiveR/HiveR.html). Rarefaction analysis for species, plasmids, strains and resistance genes was performed using the iNEXT R package^108^.

### Statistical Analysis

Statistical tests were carried out using R and are two-sided unless otherwise specified. For enrichment analysis at the cluster level (overlap across time, cohorts or resistance status) Fisher’s exact test was used, and for analysis at the genome level (fraction of genomes with a specific property) the binomial test was used.

### Data and source code availability

All sequencing reads and assemblies are available from the European Nucleotide Archive under project PRJEB31632 (https://www.ebi.ac.uk/ena/data/view/PRJEB31632). Source code and associated data for reproducing the figures are available on GitHub under an MIT license (https://github.com/csb5/hospital_microbiome). Assemblies and annotations for genomes, plasmids and phages from this study are also available at https://ndownloader.figshare.com/files/17290511.

## Acknowledgements

Funding for this work was provided by A*STAR, and we are grateful for support from NMRC (NMRC CGAug16C005). C.E.M. acknowledges support from the WorldQuant Foundation, the Bill and Melinda Gates Foundation (OPP1151054), and the Alfred P. Sloan Foundation (G-2015-13964). The funders had no role in study design, data collection and analysis, decision to publish, or preparation of the manuscript. We would like to thank Prof Jack Gilbert for insightful comments and feedback on this work. We would also like to acknowledge constructive feedback and valuable guidance from the reviewers, particularly in accounting for laboratory contaminants and artefacts.

