## Supplementary FIgures for "Cartography of opportunistic pathogens and antibiotic resistance genes in a tertiary hospital environment"

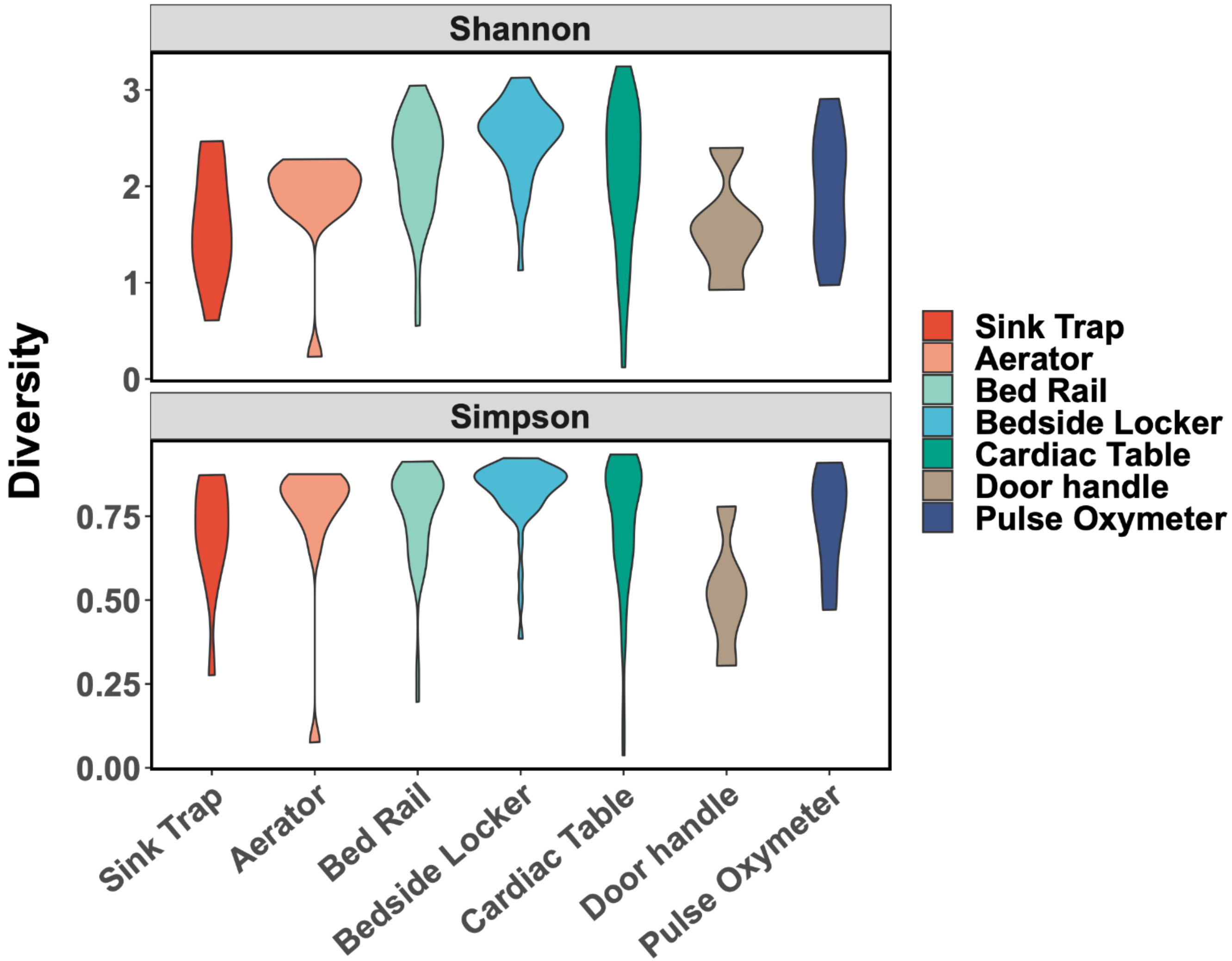


**Supplementary Figure 1:** Violin plots showing the distribution of genus-level (top) Shannon and (bottom) Simpson diversity metrics for different sampled sites. Shannon diversity of CTA sites was generally higher than CTB sites (Wilcoxon p-value<$10^{-3}$).

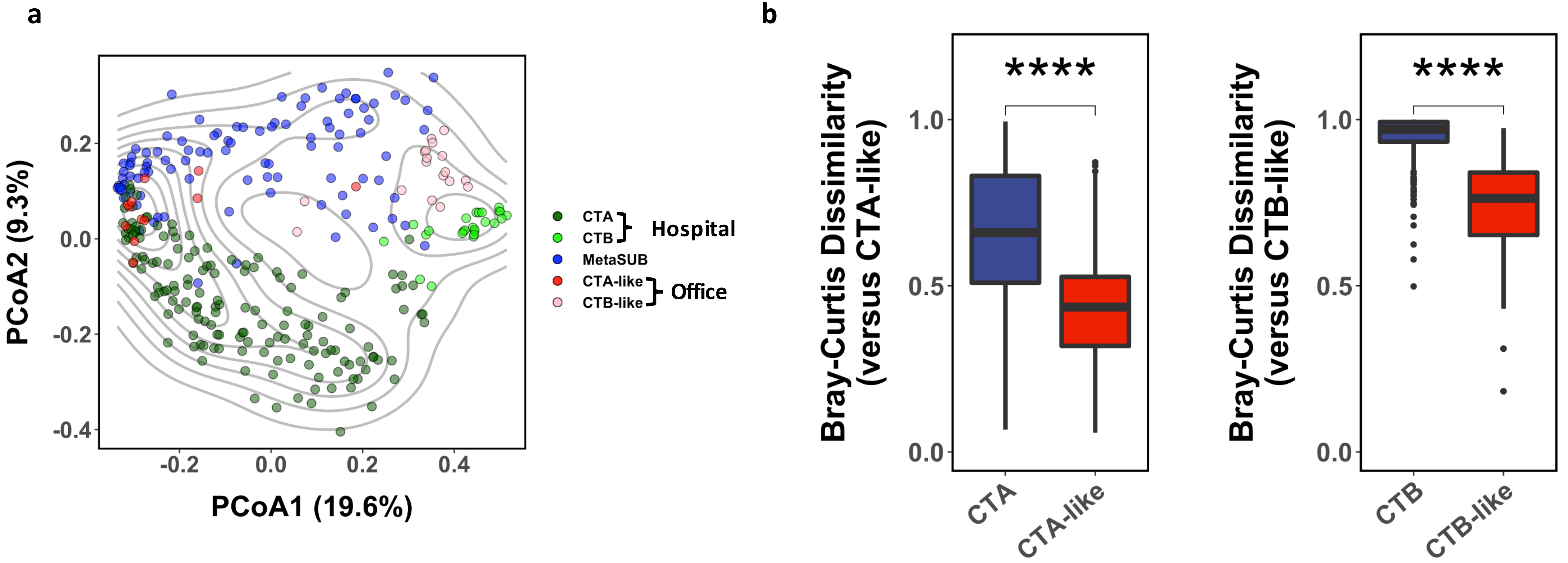


**Supplementary Figure 2:** a) Principle coordinates analysis plot (genus-level Bray-Curtis dissimilarity) based on taxonomic profiles for hospital, office and other high-touch environmental microbiomes (MetaSUB; Singapore samples). b) Boxplots showing that hospital CTA and CTB microbiomes are distinct from corresponding office microbiomes (CTA-like: office desk, chair handle, door handle, keyboard; CTB-like: sink trap, aerator; genus-level Bray-Curtis dissimilarity, **** = Wilcoxon p-value<0.0001). Boxplots are represented with center line: median; box limits: upper and lower quartiles; whiskers: 1.5× interquartile range; points: outliers.

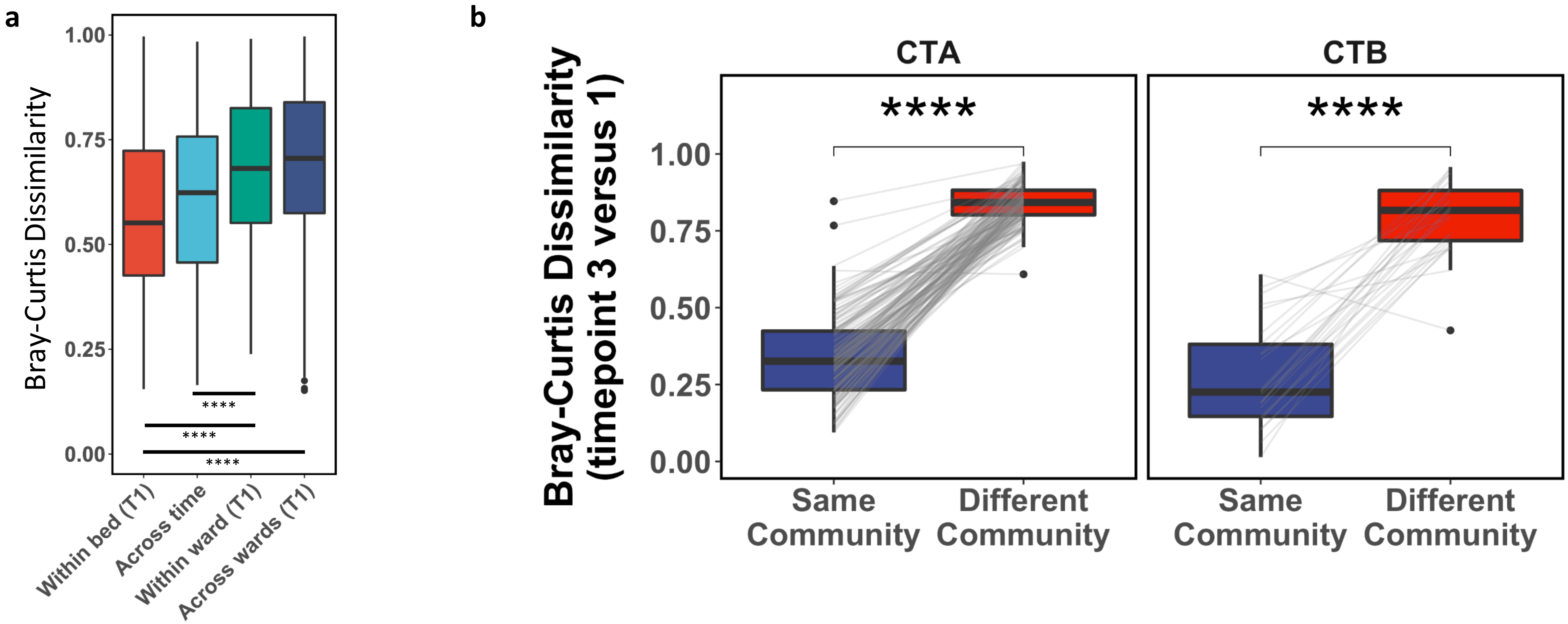


**Supplementary Figure 3:** a) Boxplots showing how dissimilarity between microbiomes (genus-level Bray-Curtis) varies as we move from sites surrounding the same bed at one timepoint (cardiac table, bed rail and bedside locker; timepoint 1), to sites associated with the same bed across two timepoints one week apart, to sites in the same ward at one timepoint (timepoint 1), and finally to those in different wards (timepoint 1). Dissimilarities in these sites increases significantly with physical distance (****=Wilcoxon p-value<0.0001). b) Boxplots showing that community type identity is largely preserved from the 1st to 3rd timepoint (1.5 years apart), where community type A (left) or B (right) samples from the 3rd timepoint are much more similar to the same community type samples in the 1st timepoint (minimum genus-level Bray-Curtis dissimilarity, ****=Wilcoxon p-value<0.0001). Boxplots are represented with center line: median; box limits: upper and lower quartiles; whiskers: 1.5× interquartile range; points: outliers.

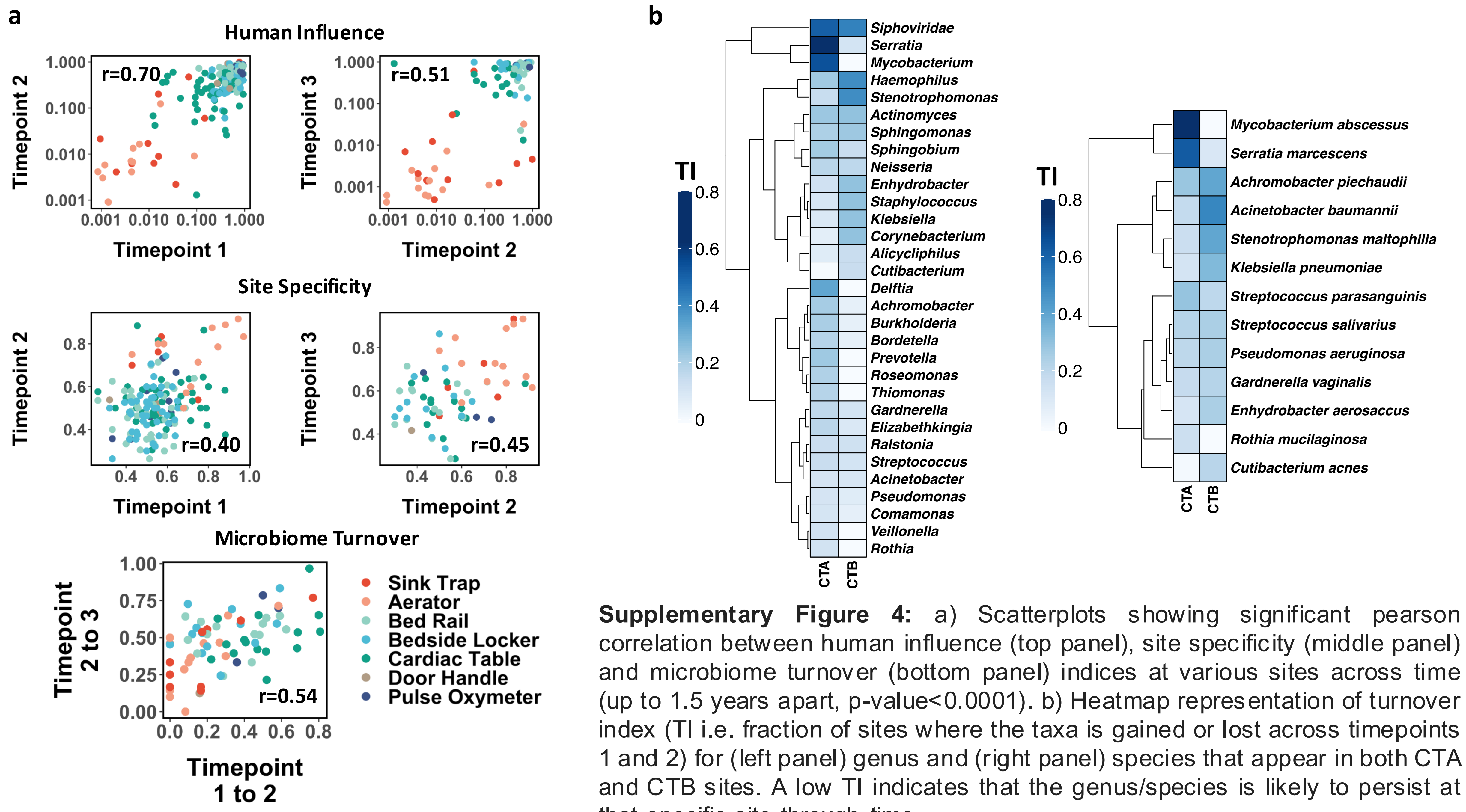


**Supplementary Figure 4:** a) Scatterplots showing significant pearson correlation between human influence (top panel), site specificity (middle panel) and microbiome turnover (bottom panel) indices at various sites across time (up to 1.5 years apart, p-value<0.0001). b) Heatmap representation of turnover index (TI i.e. fraction of sites where the taxa is gained or lost across timepoints 1 and 2) for (left panel) genus and (right panel) species that appear in both CTA and CTB sites. A low TI indicates that the genus/species is likely to persist at that specific site through time.

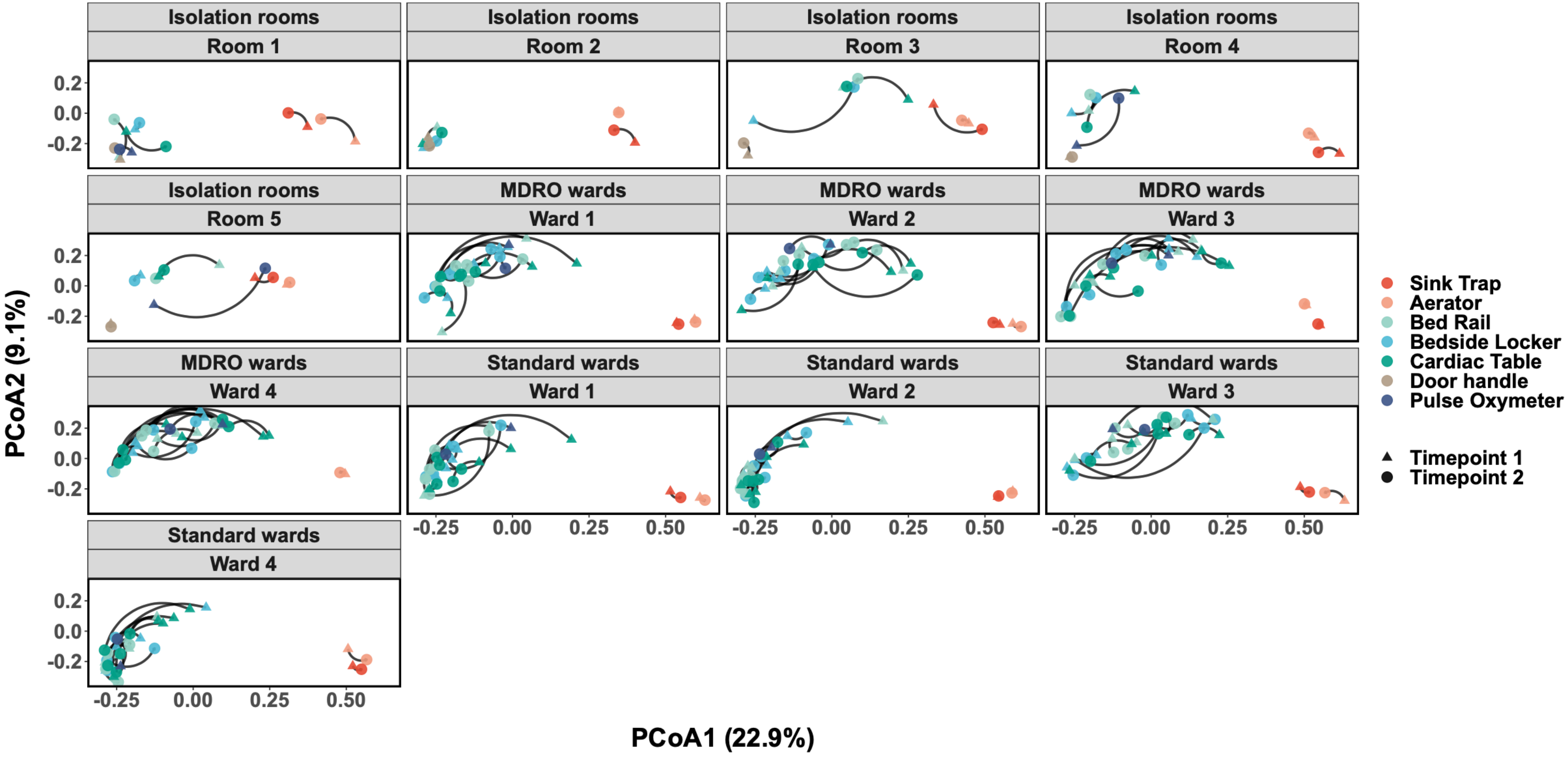


**Supplementary Figure 5:** Principle coordinates analysis (genus-level Bray-Curtis dissimilarity) of environmental microbiomes in different wards of the hospital (lines connect the same site across the two timepoints). Interestingly, CTB sites resemble CTA sites more in isolation rooms 3 and 5, while the figure further highlights the stability of CTB sites in general.

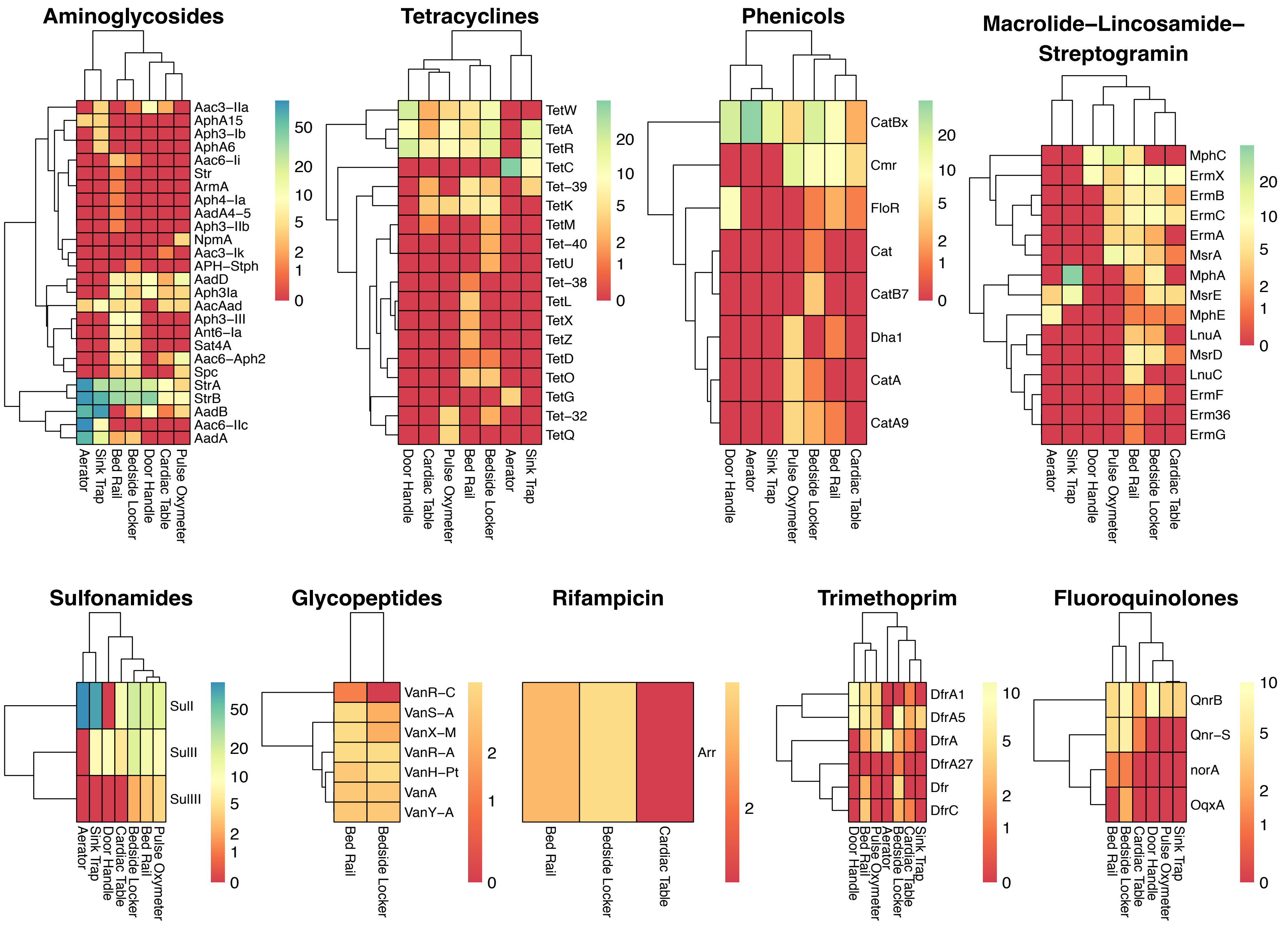


**Supplementary Figure 6:** Heatmaps showing the frequencies (percentage) at which antibiotic resistance genes were detected across sites. Zero values are depicted in red.

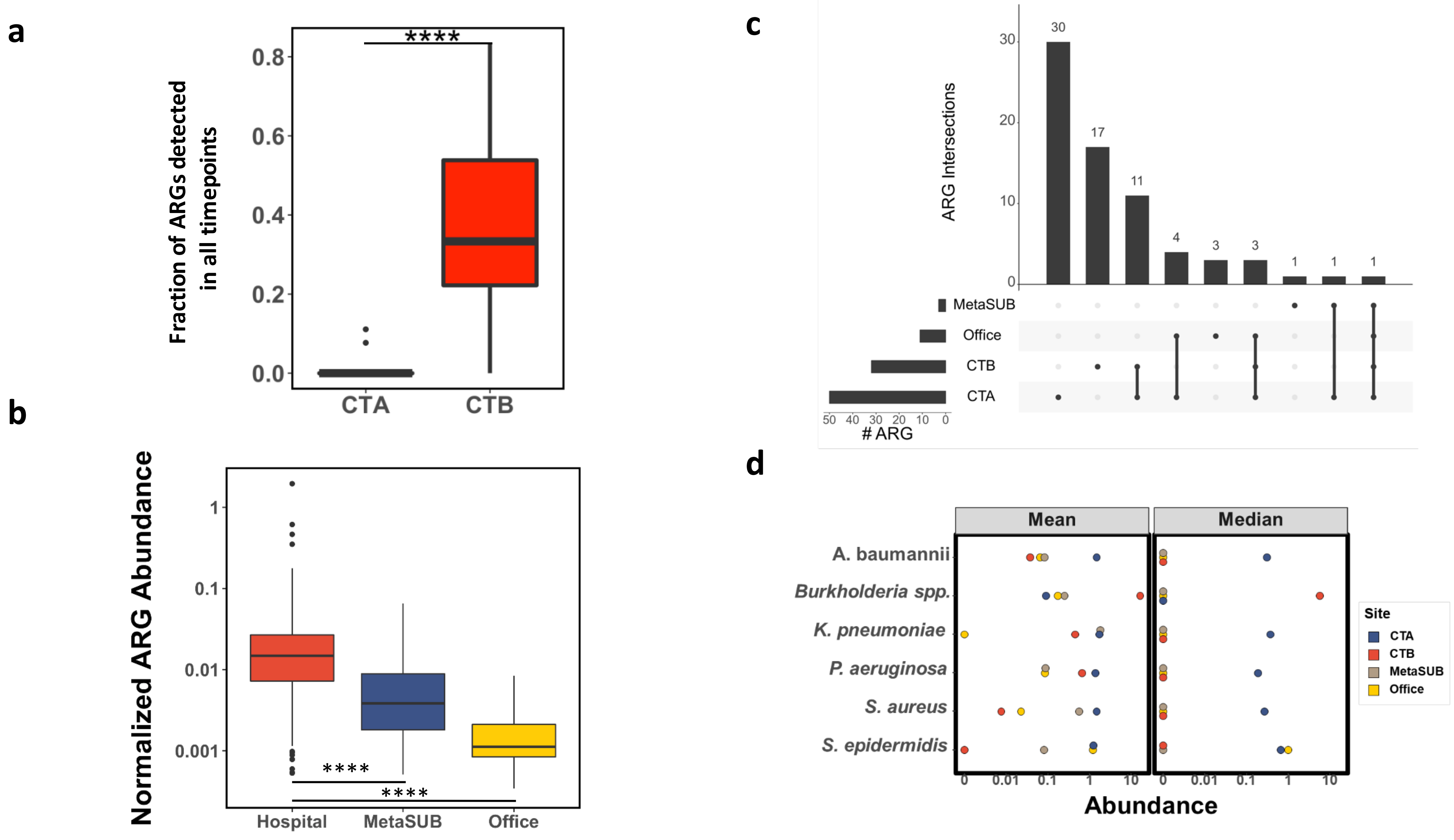


**Supplementary Figure 7:** a) Boxplots highlighting the stability of antibiotic resistance genes (ARGs) in CTB vs CTA sites. b) Boxplots showing the dramatic enrichment of ARGs in hospital environments (y-axis on log-scale). c) Upset plot showing overlaps in ARGs present in hospital (CTA or CTB sites), office and other environmental (MetaSUB; Singapore samples) microbiomes (normalized for sample size by subsampling; average of 100 replicates). d) Dotplots showing mean and median abundances of common nosocomial pathogens in the environment microbiomes of hospital (CTA or CTB), office and other community (MetaSUB) areas. Boxplots are represented with center line: median; box limits: upper and lower quartiles; whiskers: 1.5× interquartile range; points: outliers. **** = Wilcoxon p-value<0.0001.

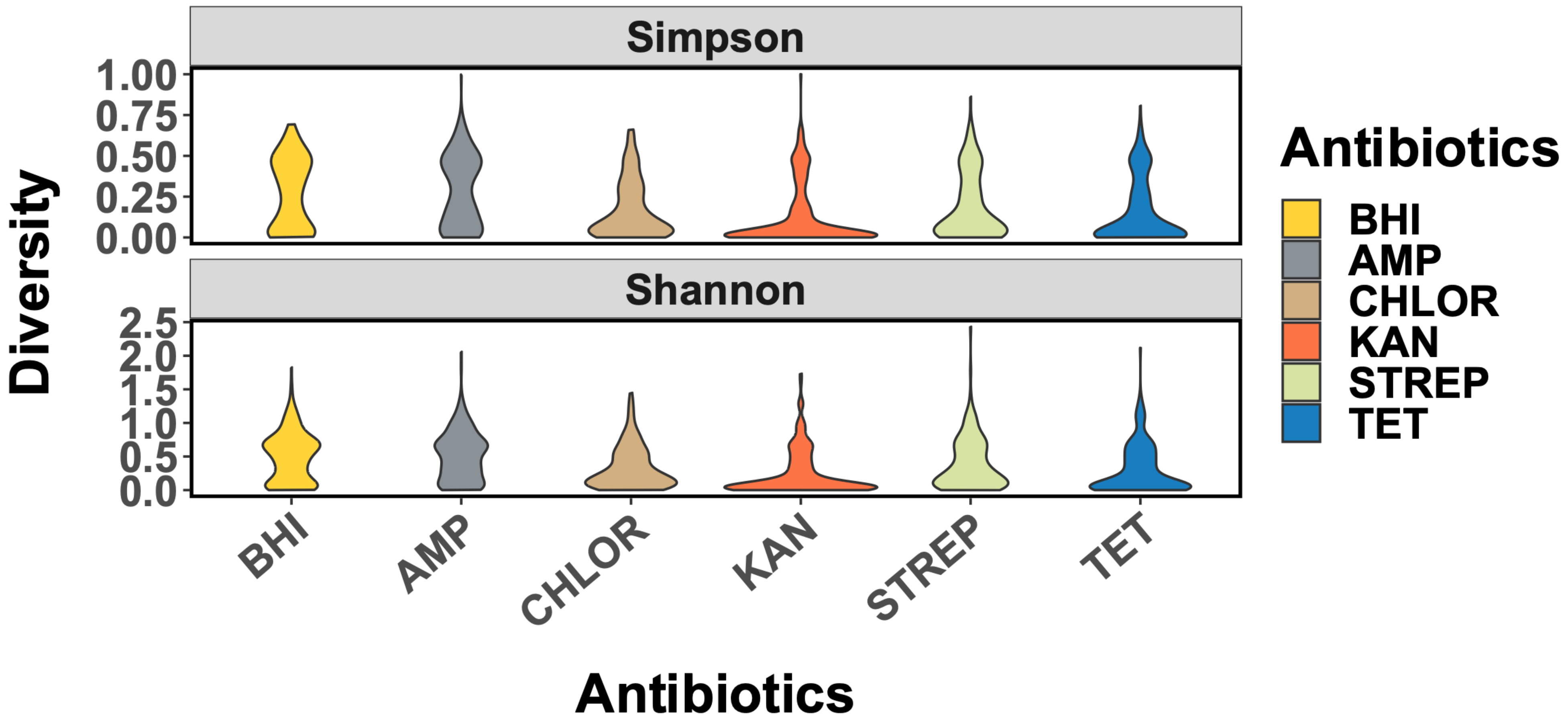


**Supplementary Figure 8:** Violin plots showing the distribution of genus-level diversity metrics for various culture-enriched communities (with BHI alone or BHI media supplemented with various antibiotics).

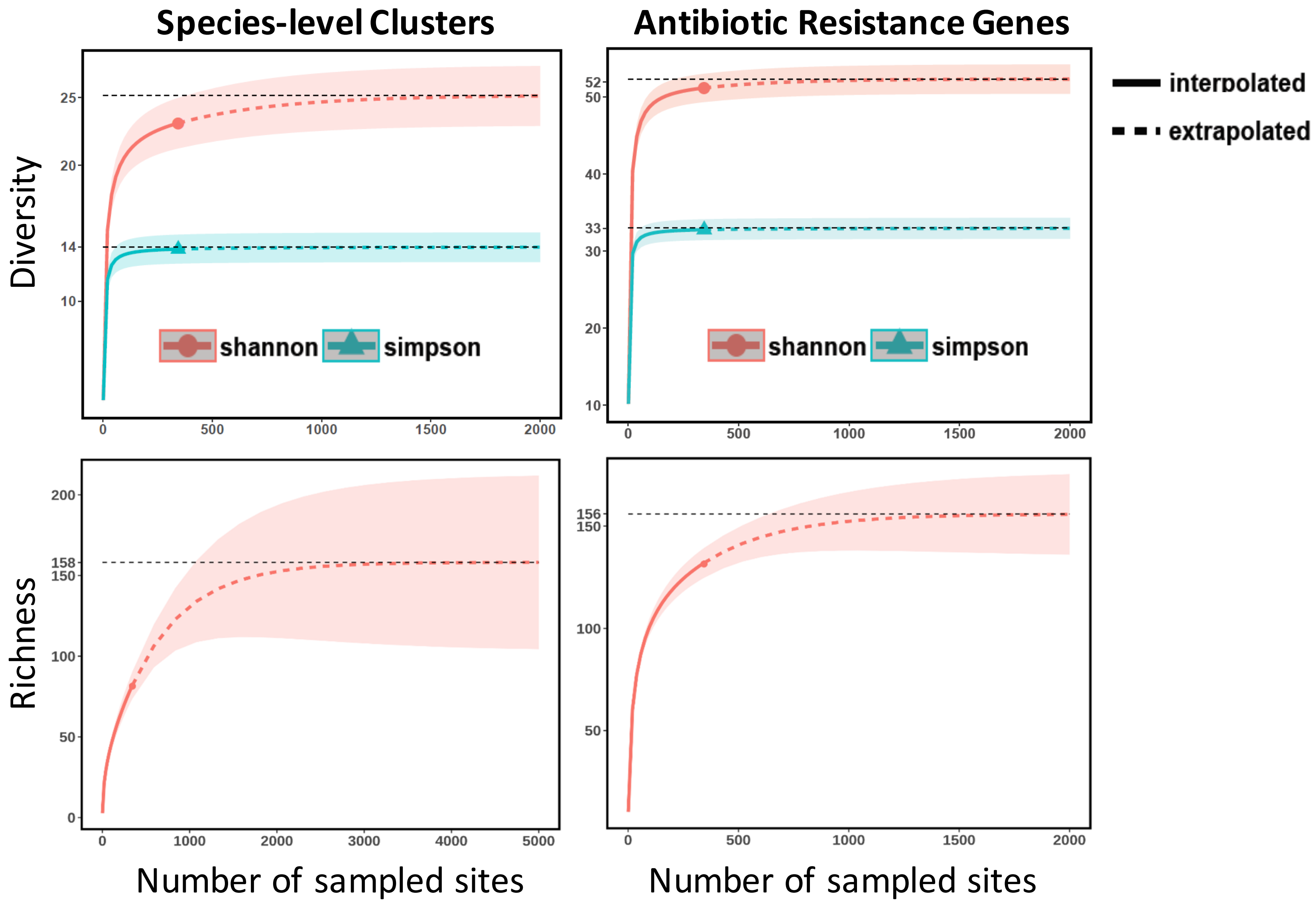


**Supplementary Figure 9:** Rarefaction analysis showing diversity (upper panel) and richness (lower panel) of species-level genomic clusters (ANI 95%) and antibiotic resistance genes observed in our genomic database as a function of the number of sites sampled. Current sampling efforts (triangle or circle) appear to capture >90% of the species and resistance gene diversity (>50% of richness) that can be sampled using this approach from the hospital environment microbiome. Shaded areas indicate 95% confidence intervals.

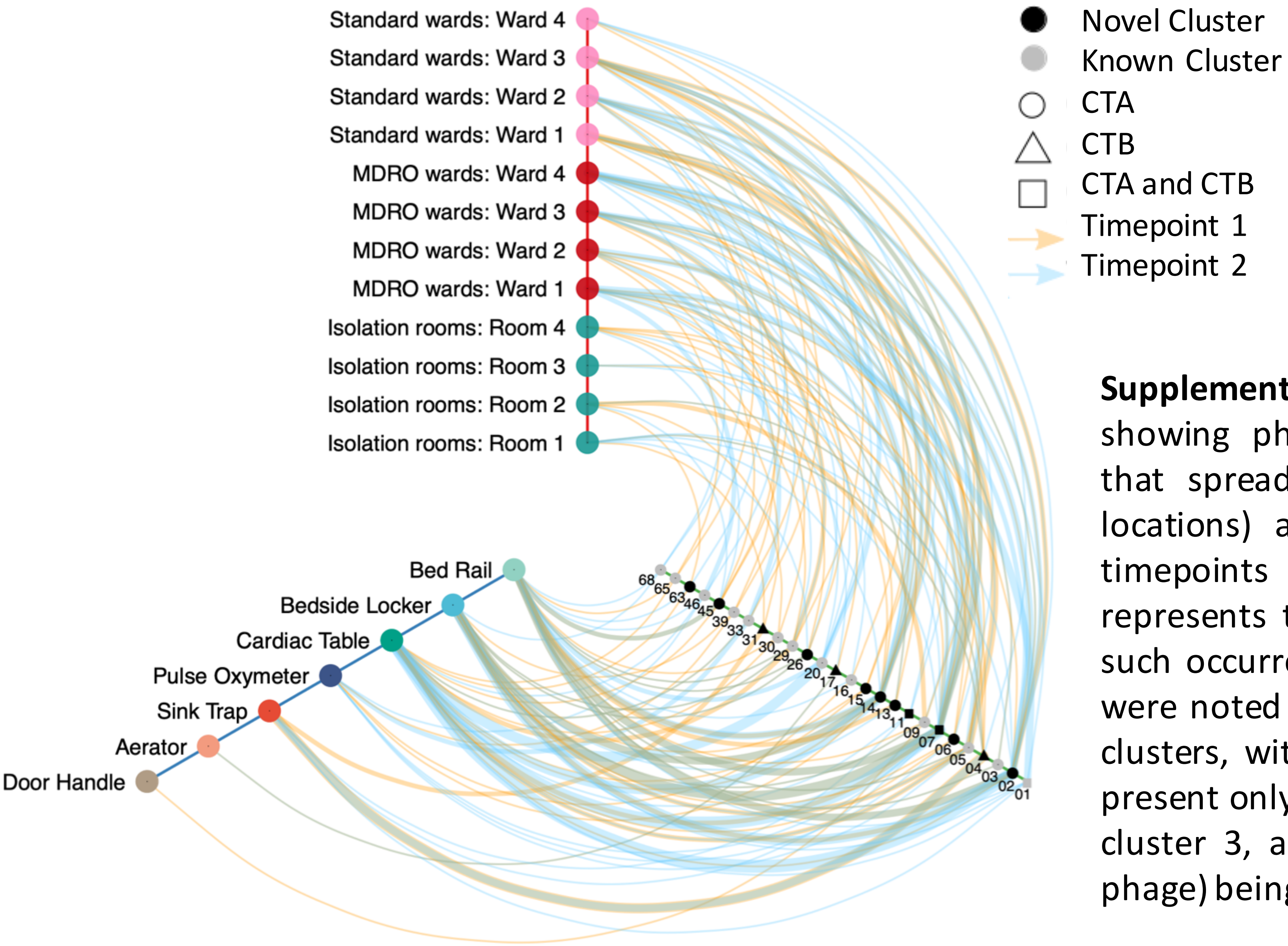


**Supplementary Figure 10**: Hive plot showing phage clusters (>99.9% ANI) that spread (observed at 2 or more locations) and/or persist (detected in timepoints 1 and 2). Line thickness represents the number of instances of such occurrences. Site-specific patterns were noted in the distribution of phage clusters, with most (77%, 20/26) being present only in CTA sites, and three (e.g. cluster 3, a novel telomere temperate phage) being present only in CTB sites.

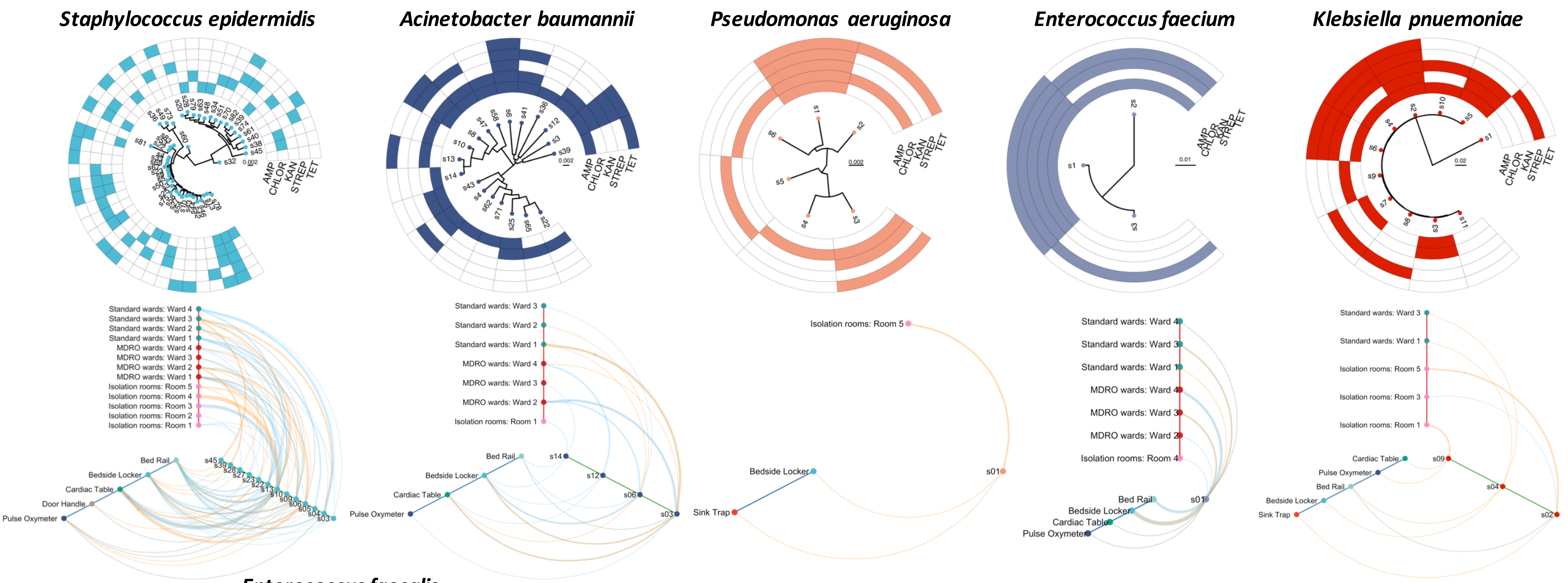


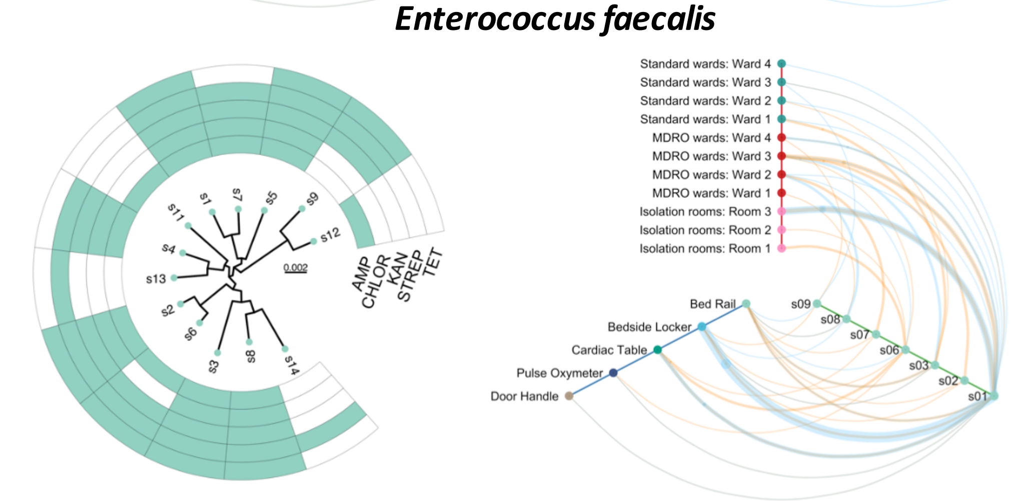


**Supplementary Figure 11:** Strain-level phylogeny (each leaf represents consensus genome of the cluster) of common nosocomial pathogens that were detected in the hospital environment with corresponding antibiotic resistance profiles, together with hive-map representation showing location of strains that spread (detected at 2 or more locations) and/or persist (detected at timepoints 1 and 2) in the hospital environment. The scale in each tree represents the number of substitutions per site, with respect to the core alignment. Orange lines represent occurrences at timepoint 1 while blue lines represent occurrences at timepoint 2. Line thickness represents the number of such observations that were made.

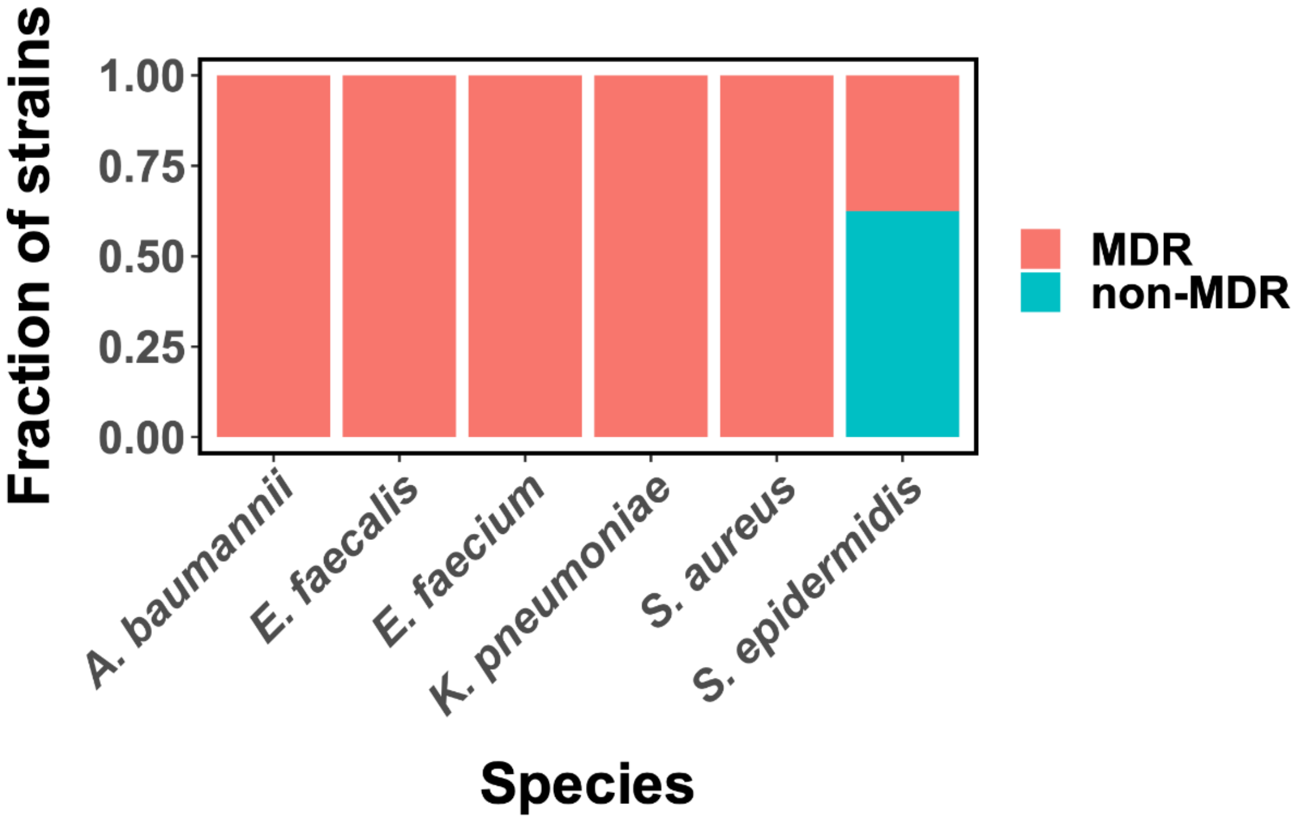


**Supplementary Figure 12:** Barplots showing the proportion of persistent (present in timepoints 1 and 2) strains that are multi-drug resistant (>2 antibiotics, MDR) for the different species.

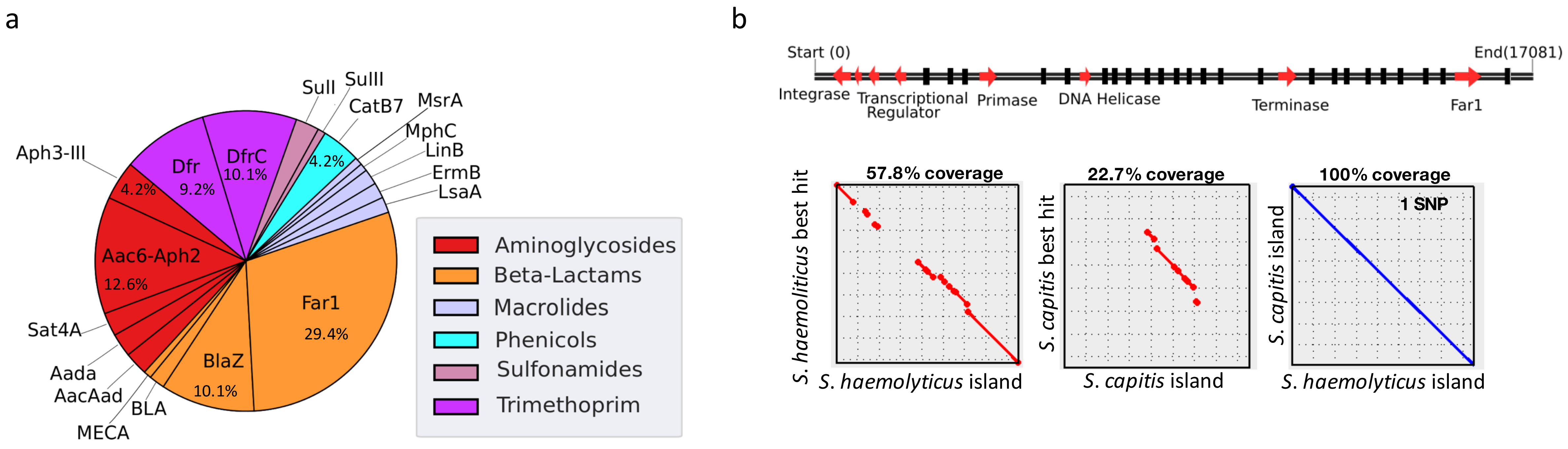


**Supplementary 13:** (a) Pie chart showing the breakdown of antibiotic resistance genes in phages/prophages/phage-like-elements. Beta-lactam resistance genes were the dominant resistance class observed (42%) with Far1 being the most common resistance gene in this class (70%). (b) (Top panel) Genome organization of a representative novel pathogenicity island from a phage-like element harboring the Far1 gene, observed in near identical copies in *S. haemolyticus* and *S. capitis* strains in the hospital environment (100% alignment, 99.994% ANI). Black lines represent hypothetical proteins and phage proteins. (Bottom Left and Center panel) Dotplots showing partial alignment between the novel pathogenicity island and its best blast hits (NCBI nt database) from *S. haemolyticus* and *S. capitis* respectively. (Bottom Right panel) Dotplot showing complete alignment between two representative pathogenicity islands found in *S. haemolyticus* and *S. capitis* cultured from the hospital environment, providing evidence for a transmission event mediated by a phage.
