## Supplementary Text for "Cartography of opportunistic pathogens and antibiotic resistance genes in a tertiary hospital environment"

### Supplementary Note 1: Assessing the impact of DNA contaminants on taxonomic profiles and identification of likely contaminant species

Following MetaSUB sample collection protocols[1], blank swabs exposed to air were collected in the hospital (handling controls) and in the laboratory environment where the samples were processed (laboratory controls). The amount of DNA extracted from handling controls was below detection limits for all swabs and hence DNA was pooled into 4 sets (from 4 samples each) for library preparation and sequencing. Comparison of hospital environment microbiomes with handling controls revealed that the taxonomic profiles observed in real samples were clearly distinct (across a range of biomass values), indicating that the impact of sampling and kitome contamination[2] on taxonomic profiles was limited (**Suppl. Note Fig. 1a**). This was further confirmed by sequencing of laboratory controls with spike-ins (*E. coli* cells and a Zymo mock community) at various concentrations (covering the range of samples that were processed in this study), where the spike-in samples exhibited very different profiles compared to blank laboratory controls (**Suppl. Note Fig. 1b**). Overall, DNA concentrations seen in libraries prepared from blank swabs were 250 to 25-fold lower than the amount seen with *E. coli* and Zymo spike-ins ($3{\times}10^5$ cells), respectively.

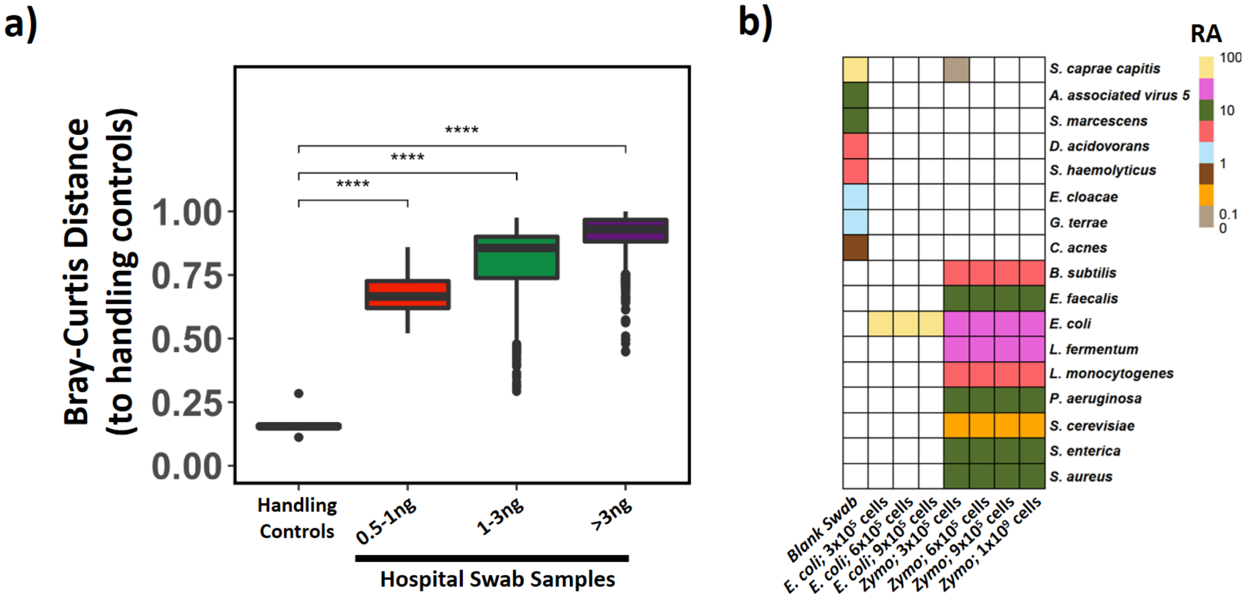


**Supplementary Note Figure 1:** a) Boxplots showing that handling controls have distinct taxonomic profiles from swabs collected in the hospital environment (genus-level Bray-Curtis dissimilarities) across a range of biomass values. More than 97% of hospital environment samples collected in this study have total DNA biomass >1ng (62% >3ng). b) Heatmap showing species-level profiles of blank swab, *E. coli* cells and a mock community (ZymoBIOMICS microbial community standard, Cat# D6300) controls for assessing the impact of the 'kitome' on taxonomic profiles of low biomass samples. Total extracted DNA biomass for the spike-ins are in parentheses: *E. coli* $3{\times}10^5$ cells (<0.1ng); $6{\times}10^5$ cells (1.2ng); $9{\times}10^5$ cells (1.8ng); mock community (Zymo) $3{\times}10^5$ cells (<0.1ng); $6{\times}10^5$ cells (<0.1ng); $9{\times}10^5$ cells (<0.1ng) and

1×10[9] cells (91ng DNA). Boxplots are represented with center line: median; box limits: upper and lower quartiles; whiskers: 1.5× interquartile range; points: outliers.

To additionally identify likely contaminant species we looked for discordance in prevalence across analysis batches[2] (timepoints 1 and 2 *versus* timepoint 3, which used different reagent kits and batches). Specifically, we identified species which were commonly present in one batch (>25% of samples with relative abundance >0.1%) but substantially less so in another (1/4$^{th}$ prevalence; red points in **Suppl. Note Fig. 2a**). The 7 species that were identified in this analysis also exhibited high correlation with each other (in 2 clusters of 5 and 2 taxa) as further evidence that they were likely contaminants[2] (**Suppl. Note Fig. 2b**). In addition, we confirmed that other potential contaminant species (close to thresholds used in **Suppl. Note Fig. 2a**) did not show high correlation with the 7 likely contaminant species (e.g. *Ruminococcus torques*, r<0.7) and/or were detected via culture based analysis (e.g. *Elizabethkingia anophelis*), and were therefore unlikely to be contaminants. A similar analysis was applied for ARGs and no genes were found to have a signature flagging them as being likely a function of laboratory contamination.

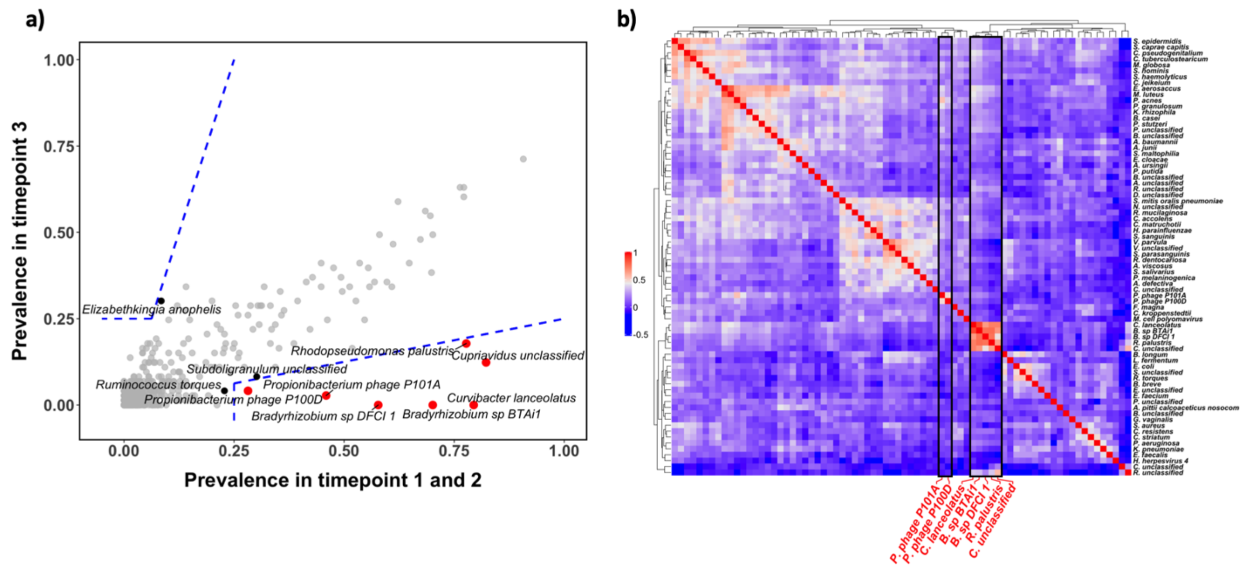


**Supplementary Note Figure 2:** a) Scatter plot showing the concordance of prevalence between batch 1 (timepoints 1 and 2) and batch 2 (timepoint 3) microbiomes. Thresholds used for identifying likely contaminant species are marked by blue lines and corresponding species are highlighted in red. Species that failed to meet the thresholds but were close (within 15%) are highlighted in black. b) Heatmap showing the correlation of abundances (Spearman) between species across samples in batch 1. Likely contaminant species are highlighted in red.

### Supplementary Note 2: Validation of culturing and antibiotic based enrichment protocols

To test for the risk of contamination during culture-based enrichment, we tested 10 blank swabs as negative controls with the same culturing protocols as used for hospital environment swabs (**Methods**). All negative controls failed to exhibit growth, even after 48 hours of incubation, suggesting that the risk of contamination from the laboratory culturing process is low. We validated the effectiveness of the culture enrichment process for selecting antibiotic resistant microbes using 6 test swabs. For each sample after culture enrichment, microbes obtained from the 5 different types of antibiotic enrichment plates (Ampicillin, Chloramphenicol, Tetracycline, Kanamycin and Streptomycin sulfate) were streaked out onto 5 separate antibiotic-free BHI agar plates. After overnight incubation at 37˚C, we picked 10 colonies from each of the antibiotic-free BHI plates (5×10 colonies in total for 1 sample) and inoculated them separately into 100 µL of BHI broth supplemented with the antibiotic that was used originally for their enrichment (Ampicillin 100 µg/mL, Chloramphenicol 35 µg/mL, Kanamycin 50 µg/mL, Streptomycin sulfate 100 µg/mL, Tetracycline 10 µg/mL). Isolates that grew (high turbidity) after incubation at 37˚C overnight helped confirm antibiotic resistance. Only 3 out of 300 isolates (1%) did not exhibit the expected antibiotic resistance (**Supplementary Note Table 1**).

| **Antibiotic Type** | **# of isolates exhibiting antibiotic resistance** | | | | | |
|---|---|---|---|---|---|---|
| | **Sample 1** | **Sample 2** | **Sample 3** | **Sample 4** | **Sample 5** | **Sample 6** |
| **Ampicillin** | 10/10 | 10/10 | 10/10 | 10/10 | 10/10 | 10/10 |
| **Chloramphenicol** | 10/10 | 10/10 | 10/10 | 10/10 | 10/10 | 10/10 |
| **Kanamycin** | 9/10 | 10/10 | 10/10 | 10/10 | 10/10 | 10/10 |
| **Streptomycin** | 9/10 | 10/10 | 10/10 | 10/10 | 10/10 | 10/10 |
| **Tetracycline** | 10/10 | 10/10 | 10/10 | 10/10 | 10/10 | 9/10 |

**Supplementary Note Table 1: Statistics for analysis confirming antibiotic resistance of isolates obtained from mixed cultures enriched with an antibiotic.**

### Supplementary Note 3: Rarefaction analysis for plasmids and strains

Rarefaction analysis for plasmids in our genomic database was used to estimate the overall diversity and richness that could have been captured. This analysis suggests that our current sampling captured >50% of the plasmid diversity (Shannon; clustered at 99% identity; 24% of richness) and a 10-fold increase in sampling (~4,000 samples) would be needed to capture the full diversity (**Supplementary Note Fig. 3a)**. Restricting the analysis to plasmids carrying antibiotic resistance genes improved sampling coverage only slightly (59% for diversity, **Supplementary Note Fig. 3b**), despite an almost complete sampling of resistance gene diversity (**Suppl. Fig. S9**). This is expected as plasmid genes can be highly mobile[3,4], consistent with the high diversity and plasticity of resistance gene combinations observed in our analysis (**Fig. 4**).

Rarefaction analysis of microbial strains also indicated that while our sampling was sufficient to reflect a majority of the strain diversity, an 8-fold increase in the size of the survey may be needed to get all strains of common nosocomial pathogens (**Supplementary Note Fig. 3c)**.

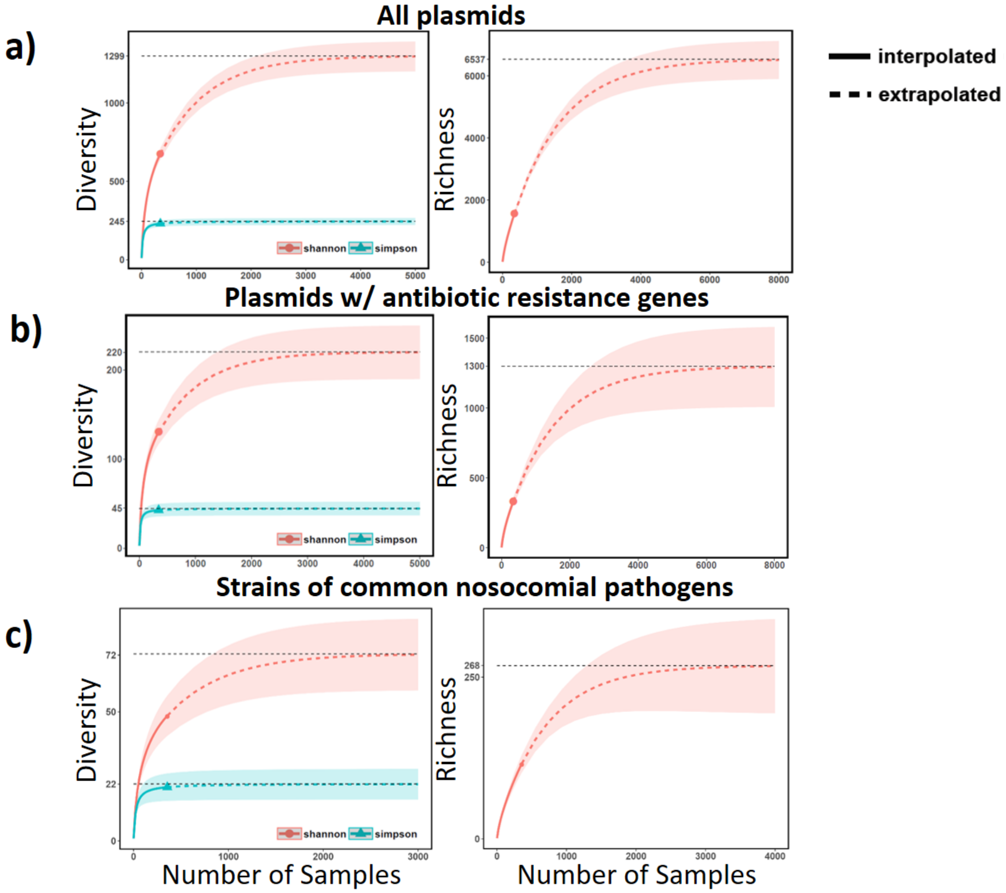


**Supplementary Note Figure 3: Rarefaction analysis for plasmids and strains in the hospital environment.** Shaded areas indicate 95% confidence intervals.
